# The mechanism for polar localization of the type IVa pilus machine

**DOI:** 10.1101/2023.06.22.546063

**Authors:** Marco Herfurth, María Pérez-Burgos, Lotte Søgaard-Andersen

**Author notes:** contributed equally to this work (Marco Herfurth, María Pérez-Burgos). Author order was determined alphabetically.

## Abstract

Type IVa pili (T4aP) are important for bacterial motility, adhesion, biofilm formation and virulence. This versatility is based on their cycles of extension, adhesion, and retraction. The conserved T4aP machine (T4aPM) drives these cycles, however the piliation pattern varies between species. To understand how these patterns are established, we focused on the T4aPM in *Myxococcus xanthus* that assembles following an outside-in pathway, starting with the polar incorporation of the PilQ secretin forming a multimeric T4aP conduit in the outer membrane. We demonstrate that PilQ recruitment to the nascent poles initiates during cytokinesis, but most is recruited to the new poles in the daughters after completion of cytokinesis. This recruitment depends on the peptidoglycan-binding AMIN domains in PilQ. Moreover, the pilotin Tgl stimulates PilQ multimerization in the outer membrane, is transiently recruited to the nascent and new poles in a PilQ-dependent manner, and dissociates after completion of secretin assembly. Altogether, our data support a model whereby PilQ polar recruitment and multimerization occur in two steps: The PilQ AMIN domains bind septal and polar peptidoglycan, thereby enabling polar Tgl localization, which then stimulates secretin multimerization in the outer membrane. Using computational analyses, we provide evidence for a conserved mechanism of T4aPM pilotins whereby the pilotin transiently interacts with the unfolded β-lip, i.e. the region that eventually inserts into the outer membrane, of the secretin monomer. Finally, we suggest that the presence/absence of AMIN domain(s) in T4aPM secretins determines the different T4aPM localization patterns across bacteria.

**Importance:** Type IVa pili (T4aP) are widespread bacterial cell surface structures with important functions in motility, surface adhesion, biofilm formation and virulence. Different bacteria have adapted different piliation patterns. To address how these patterns are established, we focused on the bipolar localization of the T4aP machine in the model organism *M. xanthus* by studying the localization of the PilQ secretin, the first component of this machine that assembles at the poles. Based on experiments using a combination of fluorescence microscopy, biochemistry and computational structural analysis, we propose that PilQ, and specifically its AMIN domains, binds septal and polar peptidoglycan, thereby enabling polar Tgl localization, which then stimulates PilQ multimerization in the outer membrane. We also propose that the presence and absence of AMIN domains in T4aP secretins determine the different piliation patterns across bacteria.

## Introduction

In bacteria, motility is important for a wide range of processes, including virulence, colonization of habitats, and biofilm formation (1, 2). Two large non-homologous envelope-spanning machines drive the two most common bacterial motility mechanisms, i.e. the extension/retraction of surface-exposed type IVa pili (T4aP) that enable cells to translocate across solid surfaces and the rotation of surface-exposed flagella that enable cells to swim through liquids or swarm across semisolid surfaces (2). Interestingly, the patterns in which these machines are positioned in cells vary between species (2–4). For both flagella and T4aP, these distinct patterns are important for efficient motility, biofilm formation and virulence (3, 5, 6). How these patterns are established is poorly understood.

T4aP are highly versatile and not only important for motility but also for surface sensing, adhesion to and colonization of host cells and abiotic surfaces, biofilm formation, virulence, predation, and DNA uptake (4, 7). The versatility of T4aP is based on their cycles of extension, surface adhesion, and retraction that are driven by the T4aP machine (T4aPM), a multiprotein complex that consists of at least 15 different proteins and spans from the outer membrane (OM) to the cytoplasm (Fig. 1) (4, 8–12). Cryo-electron tomography of the piliated and non-piliated forms of the T4aPM of *Myxococcus xanthus* and *Thermus thermophilus* revealed that both forms are multilayered structures (8, 9) (Fig. 1). However, while the architecture of the T4aPM is conserved, bacteria have adapted different piliation patterns. Specifically, in the rod-shaped cells of *Pseudomonas aeruginosa* (13, 14) and *Myxococcus xanthus* (15, 16), T4aP localization alternates between the two cell poles, while in the rod-shaped *Thermosynechococcus vulcanus* cells, they localize at both cell poles simultaneously (17). They localize in a “line along the long cell axis” (from hereon, lateral pattern) in the coccobacillus-shaped *Acinetobacter baylyi* cells (6), to the junctions between cells in the hormogonium of *Nostoc punctiforme* (18), and peritrichously in the rods of *Burkholderia cepacia* (19) and in the coccoid-shaped cells of *Neisseria meningitidis* (20)*, Neisseria gonorrhoeae* (21)*, Moraxella catarrhalis* (22) and *Synechocystis* sp. PCC6803 (23). Accordingly, the T4aPM has specifically been shown to localize to both poles in *P. aeruginosa* (14, 24, 25) and *M. xanthus* (26–29), laterally in *A. baylyi* (6), and to the intercellular junctions in *N. punctiforme* (18). To address how and when these T4aPM localization and piliation patterns are established, we focused on its bipolar localization in the model organism *M. xanthus*.

**Figure 1.**
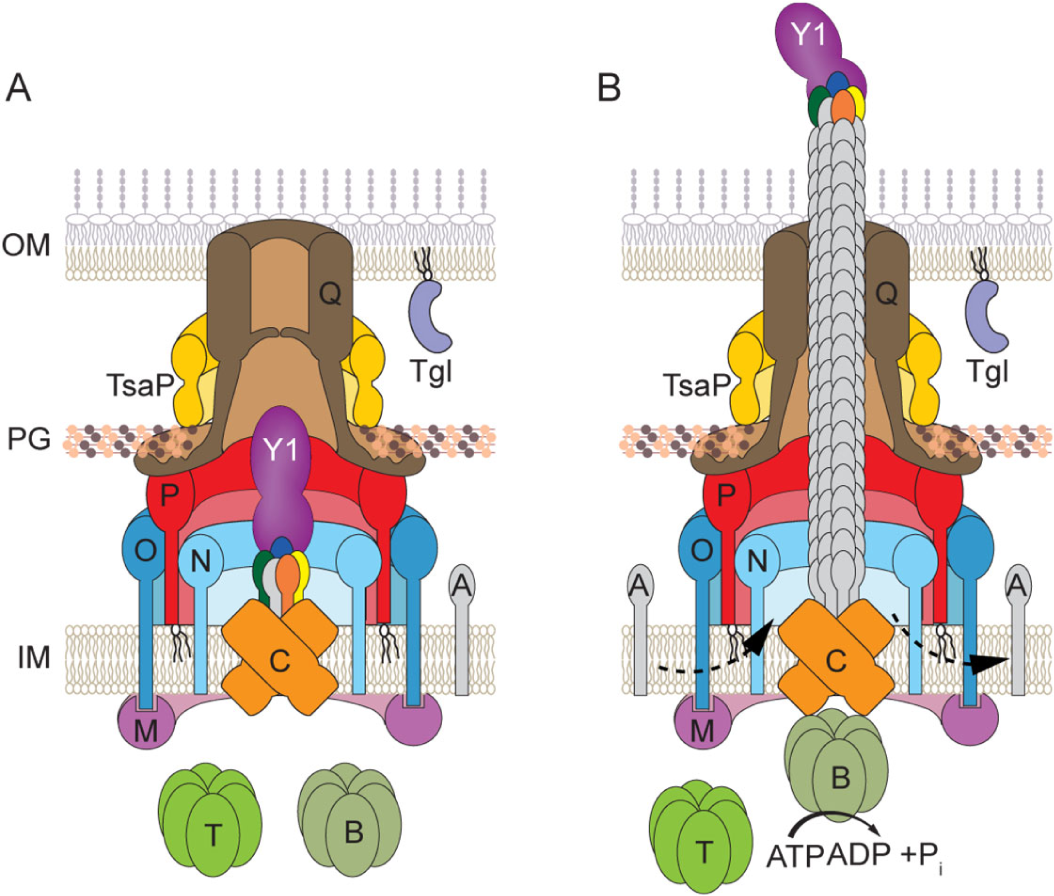
Architectural model of the T4aPM Architectural model of the cell envelope-spanning non-piliated (A) and (B) piliated T4aPM (B) of *M. xanthus* with the 15 core proteins (8). The T4aPM is divided into five parts: (1) The OM secretin channel is formed by PilQ and stabilized by the LysM domain protein TsaP (28). (2) The PilN/-O/-P periplasmic alignment complex is anchored in the inner membrane (IM) and interact with PilQ. (3) The IM/cytoplasmic platform complex is composed of PilC/-M. (4) The extension/retraction ATPases PilB/-T bind to the cytoplasmic base of the T4aPM in a mutually exclusive manner (8, 95, 96). (5) The pilus fiber is formed by PilA subunits and a priming complex, composed of PilY1 and four minor pilins (blue: PilX, green: PilW, orange: PilV, yellow: FimU), that remains at the tip of the extended T4aP (8, 12). Tgl is an OM lipoprotein that is required for PilQ secretin assembly (27, 29). Bent arrows indicate incorporation of and removal from the pilus base of PilA during extension and retraction, respectively. Proteins labeled with single letters have the Pil prefix.

As noted, *M. xanthus* assembles T4aPM at both poles, but the T4aPM are only active at one pole at a time and this pole changes on average every 10-15 min (30, 31). The assembly of the T4aPM at the two poles in *M. xanthus* depends on the OM secretin PilQ (Fig. 1) and follows an outside-in pathway (8, 27). Without the secretin, the remaining components either do not accumulate or are not incorporated into the T4aPM (27). Moreover, assembly of the T4aPM was suggested to occur at the nascent and new cell pole during and immediately after the completion of cytokinesis (27). The secretin forms the conduit for the T4aP in the OM (8, 9) (Fig. 1). In contrast to canonical OM β-barrel proteins in which a single polypeptide forms the β-barrel, the β-barrel formed by a secretin is generated from 12-15 subunits and most of the secretin pore is periplasmic (8, 32–36). Secretin protomers comprise two major subdomains, an N-terminal species-specific region and the C-terminal conserved pore-forming region (32). The N-terminal region contains at least two N-domains and, in the case of T4aPM secretins, also occasionally one or more peptidoglycan (PG)-binding AMIN domains (24, 32, 34). The C-terminal region comprises the secretin domain and the β-lip region and forms most of the barrel and a gate that closes the pore (34, 37). The periplasmic part of the secretin oligomer forms a large vestibule, which is open towards the periplasm and closed towards the OM by the gate (32). Secretins also facilitate substrate translocation across the OM in diverse other T4P systems and in type II secretion systems (T2SS) as well as type III secretion systems (T3SS) (32, 33). For their assembly in the OM, secretins rely on a cognate pilotin protein that assists in (1) secretin monomer transport to OM (38–40), (2) secretin insertion in the OM (37, 41), (3) secretin oligomerization in the OM (27, 29, 37, 39, 42), and/or (4) protection of the secretin monomer from proteolytic degradation (43, 44). Pilotins are OM periplasmic lipoproteins (33). In *M. xanthus,* the OM lipoprotein Tgl is the cognate PilQ pilotin (45–48). Lack of Tgl causes a defect in PilQ multimerization (27, 29). Consequently, in the absence of Tgl, the remaining parts of the T4aPM do not accumulate or not assemble (27). Interestingly, the lack of PilQ assembly in Δ*tgl* cells can be extracellularly complemented by Tgl^+^ cells in a process referred to as OM-exchange, in which OM proteins are exchanged between cells (45, 48–50).

To understand how T4aPM becomes polarly localized in *M. xanthus*, we investigated when and how the PilQ secretin is recruited to the poles. We show that PilQ recruitment to the nascent pole initiates during cytokinesis, but most is recruited to the new poles in the daughter cells after completion of cytokinesis. We also demonstrate that Tgl is transiently recruited to the nascent and new poles during and after cytokinesis in a PilQ-dependent manner, and that Tgl dissociates after secretin assembly is completed. Based on a dissection of PilQ, our data support that its N-terminal PG-binding AMIN domains are crucial for its septal and polar recruitment likely via binding to PG specific to the septum and cell poles. Our data support a model whereby PilQ monomers are recruited to the nascent and new cell poles by specific septal and polar PG via their AMIN domains, thereby enabling Tgl localization, and, consequently, secretin assembly in the OM. We also propose that the presence/absence of PG-binding AMIN domain(s) in T4aPM secretins is responsible for the different localization patterns of T4aPM across bacteria.

## Results

### The secretin PilQ is stably recruited to the nascent and new poles

Previously, PilQ was suggested to be recruited to the nascent and new poles of *M. xanthus* cells during and immediately after completion of cytokinesis (27). In those experiments, a partially active PilQ-sfGFP fusion that accumulated at a reduced level was used (27). To reassess PilQ recruitment to the nascent pole, we used a strain, in which an active PilQ-sfGFP fusion protein was expressed from the native site (5) (Fig. S1A). In immunoblots, the heat- and SDS-resistant PilQ multimer accumulated at close to native levels while the monomer was only detected at a very low level (Fig. 2A). Of note, a small fraction of PilQ-sfGFP was cleaved to PilQ and sfGFP (Fig. 2A).

**Figure 2.**
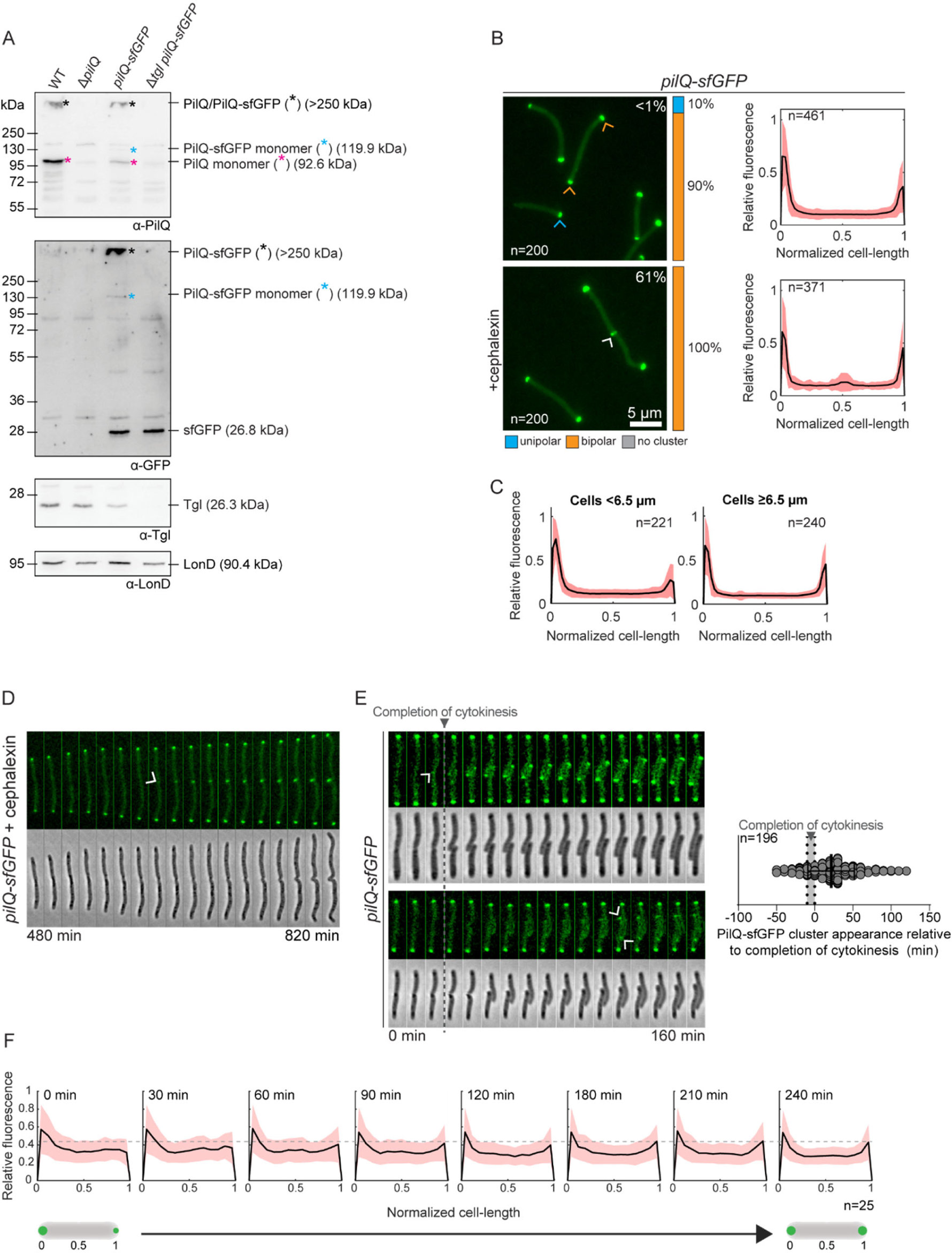
Accumulation of PilQ variants and localization of PilQ-sfGFP (A) Immunoblot detection of PilQ/PilQ-sfGFP. Protein from the same number of cells from exponentially growing suspension cultures was loaded per lane. Blot was probed with the indicated antibodies. The blot was stripped before applying a new antibody. LonD served as a loading control. Monomeric and oligomeric forms of PilQ/PilQ-sfGFP are marked with an asterisk. Calculated molecular weights of proteins without signal peptide (if relevant) are indicated. (B) Localization of PilQ-sfGFP in the presence and absence of cephalexin. Left panels, representative epifluorescence images of cells expressing PilQ-sfGFP. The percentage of cells with a unipolar (blue), bipolar (orange) cluster localization pattern or no cluster (grey) is indicated. Blue, orange and white arrowheads indicate unipolar, bipolar and mid-cell clusters. Percentage of cells with a mid-cell cluster is indicated in white. Right panels, normalized fluorescence profiles of cells, for which a cluster was detected, as a function of the relative cell length. Mean (black line) and standard deviation (SD) (orange) of the relative fluorescence along the normalized cell are depicted. Cell length was normalized from 0 to 1, where 0 is assigned to the pole with the highest fluorescent value. (C) Localization of PilQ-sfGFP in short (<6.5 µm) and long (≥6.5 µm) cells. Same cells analyzed as in untreated cells in B. Signals are shown as in B, right panel. (D) Time-lapse microscopy of a PilQ-sfGFP expressing cell treated with cephalexin. Epifluorescence and phase-contrast images are shown. Arrow indicates first time point at which the PilQ-sfGFP cluster is clearly visible at mid-cell. Time indicates time point after the addition of cephalexin (t=0). (E) Polar recruitment of PilQ-sfGFP during the cell cycle. Left panels, epifluorescence and phase-contrast images from time-lapse microscopy of cells expressing PilQ-sfGFP. PilQ-sfGFP clusters are visible at the nascent poles during (upper panel) or after completion (lower panel) of cytokinesis. Arrowheads indicate first time point at which the PilQ-sfGFP cluster is clearly visible. Right panel, analysis of appearance of the PilQ-sfGFP cluster relative to completion of cytokinesis for each daughter cell. The first time point after completion of cytokinesis is defined as t=0 and indicated by the grey vertical bar. The black line and error bars represent the mean ± SD. The appearance of PilQ-sfGFP clusters was studied in 196 daughter cells. (F) Analysis of polar incorporation of PilQ-sfGFP after completion of cytokinesis. The cellular fluorescence was quantified at different time points after completion of cytokinesis and the relative fluorescence along a normalized cell was plotted. Mean (black line) and SD (orange) are indicated. Cell length was normalized from 0 to 1, where 0 was assigned to the old pole. n=25. In D-F, to follow cells on hard agar by time-lapse microscopy for extended periods of time and avoid that they move out of the field of view, all strains contain an in-frame deletion of *gltB* (Δ*gltB),* which encodes a component of the *M. xanthus* gliding motility machine (85, 97).

In agreement with previous observations that the *M. xanthus* T4aPM assembles at both cell poles (8, 26–29), PilQ-sfGFP overall localized in a bipolar pattern (Fig. 2B) and ∼19±6% of the fluorescent signal is polar (5). However, we noticed that long cells had more symmetric bipolar PilQ-sfGFP clusters, while short cells had a higher degree of asymmetry and a few short cells even only had a unipolar signal (Fig. 2B and C). We did not reliably identify dividing cells with PilQ-sfGFP at the nascent poles at the constriction site at mid-cell.

To determine whether PilQ-sfGFP can be recruited to the nascent cell poles during cytokinesis, we treated cells with cephalexin to inhibit FtsI that catalyzes PG cross-linking at the septum (51), and blocks cytokinesis after the initiation of constriction in *M. xanthus* (52, 53). In cells treated with cephalexin for 4-5 h, largely corresponding to one doubling time, the cell length had increased, the bipolar PilQ-sfGFP signals were more symmetrical, unipolar PilQ-sfGFP localization was not observed, and, importantly, PilQ-sfGFP localized at the constriction site at mid-cell in two-thirds of the cells (Fig. 2B). The cluster at the constriction site was stable when treated cells were followed by time-lapse fluorescence microscopy (Fig. 2D). These observations demonstrate that PilQ-sfGFP can be stably incorporated into the nascent poles during cytokinesis. Consistent with these findings, we observed by time-lapse fluorescence microscopy of untreated PilQ-sfGFP-expressing cells that ∼20% of cells had a very faint PilQ-sfGFP cluster at the constriction site at mid-cell up to 50 min prior to completion of cytokinesis (Fig. 2E). However, most of the clusters only became clearly visible at the new poles after completion of cytokinesis, and on average, a polar cluster became reliably visible 20 min after completion of cytokinesis. In the daughter cells, the PilQ-sfGFP clusters at the new poles increased in intensity over time, and mostly during the first 60-90 min after completion of cytokinesis, ultimately resulting in the more symmetric bipolar localization pattern (Fig. 2E and F).

We conclude that recruitment of PilQ to the nascent poles initiates during cytokinesis but most of PilQ is recruited over the first 60-90 min after completion of cytokinesis resulting in a symmetric bipolar localization of PilQ. We speculate that we did not detect a PilQ-sfGFP signal at the site of division in the analysis of snapshots (Fig. 2B) because the PilQ-sfGFP signal before completion of cytokinesis is too faint to be reliably detected and only becomes reliably detected when cells are followed in time-lapse microscopy experiments.

### The pilotin Tgl is transiently recruited to the nascent and new poles

Next, we investigated the localization of the pilotin Tgl. We previously analyzed Tgl localization using a strain overexpressing an active Tgl-sfGFP protein and found that it localized to the cell envelope but not specifically at the cell poles or the division site (27). By contrast, Nudleman et al. found by immunostaining that Tgl localized unipolarly in ∼30% of the cells (29). To resolve the localization of Tgl, we generated a strain expressing the active Tgl-sfGFP fusion (Fig. S1A) from the native site at native levels (Fig. 3A) and reevaluated its localization.

**Figure 3.**
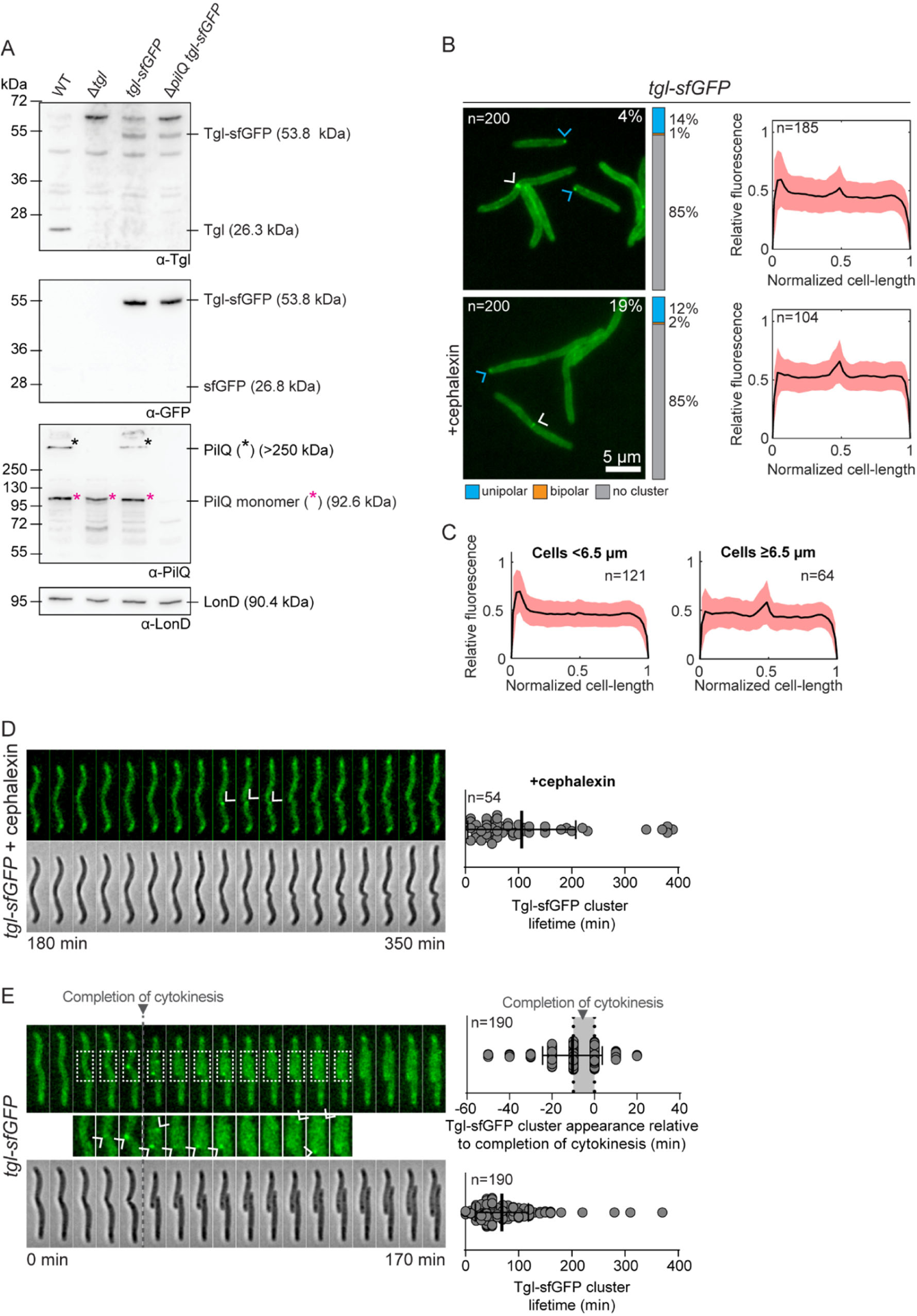
Accumulation of Tgl variants and localization of Tgl-sfGFP (A) Immunoblot detection of Tgl/Tgl-sfGFP. Protein from the same number of cells from exponentially growing suspension cultures was loaded per lane. Blot was probed with the indicated antibodies. The blot was stripped before applying a new antibody. LonD served as a loading control. Monomeric and oligomeric forms of PilQ are marked with an asterisk. Calculated molecular weights of proteins without signal peptide (if relevant) are indicated. (B) Localization of Tgl-sfGFP in the presence and absence of cephalexin as in Fig. 2B. (C) Localization of Tgl-sfGFP in short (<6.5 µm) and long (≥6.5 µm) cells. Same cells analyzed as in untreated cells in panel B. Signals are shown as in Fig. 2B, right panel. (D) Time-lapse microscopy of cells expressing Tgl-sfGFP treated with cephalexin. Left panel, epifluorescence and phase-contrast images are shown. Arrows indicate time points at which the Tgl-sfGFP cluster is clearly visible. Time indicates time point after the addition of cephalexin (t=0). Right diagram shows lifetime of Tgl-sfGFP clusters. The black line and error bars represent the mean ± SD. n=54. (E) Recruitment of Tgl-sfGFP to the nascent and new poles during the cell cycle. Left panel, epifluorescence and phase-contrast images from time-lapse microscopy of a cell expressing Tgl-sfGFP. Tgl-sfGFP clusters generally appear at the nascent poles during cytokinesis. Arrows indicate time points, at which the Tgl-sfGFP cluster is clearly visible. The boxed areas are shown below in a higher magnification. Right panels, analysis of appearance of a Tgl-sfGFP cluster relative to completion of cytokinesis for each daughter cell and lifetime of the Tgl-sfGFP cluster relative to completion of cytokinesis for each daughter cell. The first time point after completion of cytokinesis is defined as t=0 and indicated by the grey vertical bar. The black line and error bars represent the mean ± SD. The appearance of Tgl-sfGFP clusters was studied in 190 daugther cells. In D-E, strains analyzed contain the Δ*gltB* mutation.

In all cells, Tgl-sfGFP localized along the entire cell periphery in a pattern typical of proteins localizing to the cell envelope (Fig. 3B). Moreover, in 14% of the cells, Tgl-sfGFP also localized in a unipolar cluster, and these cells were typically short in length (Fig. 3B and C). Additionally, in 4% of the cells, Tgl-sfGFP localized at the constriction site, and these were typically long cells (Fig. 3B and C). In the remaining cells, Tgl-sfGFP did not form clusters (Fig. 3B). We note that Tgl-sfGFP localization is very different from the bipolar localization of the T4aPM in *M. xanthus* cells. We speculate that in our previous analysis of Tgl-sfGFP localization, its overexpression and the resulting strong cell envelope signal likely masked the weak Tgl-sfGFP clusters at the nascent and new poles.

Treatment of Tgl-sfGFP-expressing cells with cephalexin caused a significant increase in the fraction of cells with a mid-cell cluster, and while the fraction of cells with a unipolar signal remained unchanged, this signal was substantially weaker than in untreated cells (Fig. 3B). Of note, the fraction of cells with Tgl-sfGFP at mid-cell was significantly lower than in the case of PilQ-sfGFP in cephalexin-treated cells (Fig. 2B). When cephalexin-treated cells were followed by time-lapse fluorescence microscopy, we observed Tgl-sfGFP clusters appear at mid-cell in cells with constrictions, and these clusters disintegrated after ∼100 min (Fig. 3D). Similarly, time-lapse fluorescence microscopy of untreated Tgl-sfGFP-expressing cells showed that the protein on average appeared at mid-cell in constricting cells ∼10 min before completion of cytokinesis but in a few cells the cluster appeared up to 50 min prior to completion of cytokinesis (Fig. 3E). Upon completion of cytokinesis, the two daughters each inherited a cluster at the new pole that eventually disintegrated (Fig. 3E).The lifetime of a cluster from its first appearance until it permanently disintegrated was ∼70 min (Fig. 3E). Given a generation time of 5-6 h, this lifetime correlates well with the percentage of cells with unipolar and mid-cell clusters quantified in snapshots of cells expressing Tgl-sfGFP (Fig. 3B). Interestingly, the lifetime of a Tgl-sfGFP cluster coincides with the time (60-90 min) required for polar incorporation of PilQ-sfGFP at the nascent and new poles (Fig. 2E and F).

The observations that the PilQ-sfGFP cluster stably remains at mid-cell in cephalexin-treated cells (Fig. 2D), while the Tgl-sfGFP cluster disintegrates in the presence or absence of cephalexin, support that Tgl is transiently localized to the nascent and new poles to promote secretin assembly in the OM and is not part of the fully assembled T4aPM.

### Tgl is important for multimerization and stability of PilQ and PilQ is important for polar recruitment of Tgl

Our fluorescence microscopy analyses showed that Tgl-sfGFP on average formed a visible cluster at mid-cell slightly earlier than PilQ-sfGFP (Fig. 2E vs Fig. 3E). We, therefore hypothesized that Tgl could be responsible for recruiting PilQ to mid-cell during cytokinesis. To this end, we analyzed protein accumulation and localization of each fluorescent fusion in the absence of the other.

In agreement with previous observations (27, 29), only the monomer fraction of PilQ and PilQ-sfGFP accumulated in the Δ*tgl* mutant (Fig. 2A and 3A), confirming that Tgl is important for multimerization of PilQ. We also noticed that the total level of the PilQ variants, and especially of PilQ-sfGFP, was reduced in the absence of Tgl (Fig. 2A and Fig. 3A), arguing that Tgl is also important for PilQ stability. Accordingly, the PilQ-sfGFP fluorescent signal was strongly reduced, and as reported (27), polar and mid-cell clusters were not detected (Fig. 4A). In previous immunofluorescence studies using a Δ*tgl*:: *tet*^R^ strain, PilQ was reported to localize to the poles in the absence of Tgl (29), however, we did also not observe PilQ-sfGFP clusters in this strain background (Fig. S1B and C).

**Figure 4.**
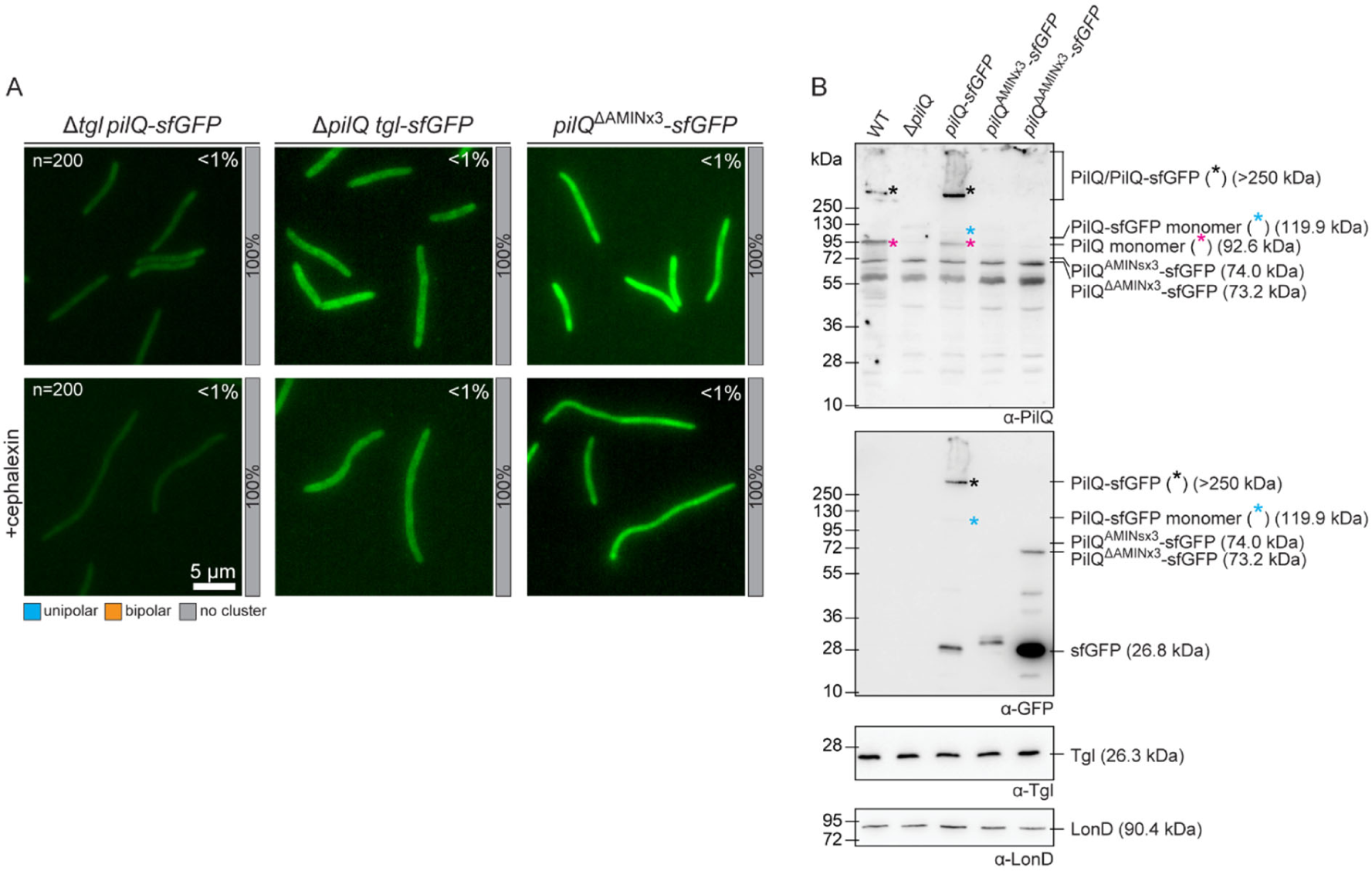
Polar Tgl-sfGFP localization depends on PilQ and polar PilQ-sfGFP localization depends on its AMIN domains. (A) Localization of PilQ-sfGFP and Tgl-sfGFP in the absence of the other as well as localization of PilQ^ΔAMINx3^-sfGFP in the presence and absence of cephalexin as in Fig. 2B, left panel. (B) Immunoblot detection of PilQ^AMIN×3^-sfGFP and PilQ^ΔAMIN×3^-sfGFP. Protein from the same number of cells from exponentially growing suspension cultures was loaded per lane. The same blot was stripped before applying a new antibody. LonD served as a loading control. Monomeric and oligomeric forms of the PilQ-sfGFP variants are marked with an asterisk. Calculated molecular weights of proteins without signal peptide (if relevant) are indicated.

In the inverse experiment, we observed that Tgl and Tgl-sfGFP accumulated at the same level in the presence and absence of PilQ (Fig. 2A and Fig. 3A) and that Tgl-sfGFP localized at the cell envelope (Fig. 4A). However, Tgl-sfGFP neither formed unipolar nor mid-cell clusters in the absence of PilQ (Fig. 4A). Because Tgl-sfGFP accumulates at native levels but does not form polar or mid-cell clusters in the absence of PilQ, these data support that PilQ recruits Tgl to mid-cell and the poles rather than the other way around. We note that the time-lapse fluorescence microscopy analyses showed that Tgl-sfGFP on average formed a visible cluster at mid-cell slightly earlier than PilQ-sfGFP (Fig. 2E vs Fig. 3E). Because cells expressing PilQ-sfGFP also accumulate a fraction of untagged PilQ monomer (Fig. 2A), we speculate that the slight delay (30 min) in the average timing of PilQ-sGFP recruitment to the nascent/new poles relative to the average recruitment of Tgl-sfGFP could originate from a preference of untagged PilQ for the constriction site.

### PilQ recruitment to the nascent and new poles depend on the AMIN domains

Next, we addressed how PilQ is recruited to the nascent poles. In *P. aeruginosa*, localization to the division site of the inner membrane (IM) protein PilO (Fig. 1), and therefore the T4aPM, to the nascent poles at the constriction site depends on the PG-binding AMIN domains of PilQ (24). The *M. xanthus* PilQ contains three AMIN domains, one of which is sufficient for the correct assembly and polar localization of the T4aPM (8). We, therefore, speculated that deletion of all three AMIN domains would prevent PilQ from being recruited to the nascent and new poles.

To this end, we generated a strain expressing a PilQ variant lacking all three AMIN domains fused to sfGFP (PilQ^ΔAMIN×3^-sfGFP) from the native site. In immunoblots with α-GFP antibodies, monomeric PilQ^ΔAMIN×3^-sfGFP accumulated at high levels, although a significant fraction of the protein was also cleaved to generate free sfGFP, and no heat-resistant multimers were detected (Fig. 4B). Consistently, cells expressing PilQ^ΔAMIN×3^-sfGFP were non-motile (Fig. S1A). Because PilQ^ΔAMIN×3^-sfGFP was not detected by the PilQ antibodies (Fig. 4B), we suggest that the epitopes detected by these antibodies are within the AMIN domains. Consistent with the three AMIN domains being essential for polar PilQ recruitment, PilQ^ΔAMIN×3^-sfGFP did not generate mid-cell and polar clusters (Fig. 4A). We conclude that the three AMIN domains are required for recruitment of PilQ to the nascent and new poles and multimer formation.

Next, to test whether the three AMIN domains are sufficient for polar recruitment, we generated a fusion in which the three AMIN domains were fused to sfGFP (PilQ^AMINs^-sfGFP). However, immunoblot analysis and fluorescence microscopy revealed that PilQ^AMINs^-sfGFP did not accumulate thus precluding further analyses (Fig. 4B and Fig. S1D).

### Cell division-independent polar recruitment of PilQ

Generally, septal and polar PG contains fewer stem peptides, is considered metabolically mostly inert and modifications acquired during cytokinesis are retained at the poles indefinitely (54, 55). Moreover, it has been shown that the AMIN domain of the cell division protein AmiC in *Escherichia coli* binds to septal PG during cytokinesis (56–58). Based on these considerations, and because the three PilQ AMIN domains are required for polar recruitment of PilQ, we hypothesized that the old cell poles would have the properties required for recruitment and incorporation of the PilQ secretin in the OM independently of a cell division event. To test this hypothesis, we expressed PilQ-sfGFP in a Δ*pilQ* mutant under the control of the vanillate-inducible promoter (P_van_) and then followed its polar recruitment (Fig. 5A and Fig. S2A). Remarkably, we observed that PilQ-sfGFP was recruited to both poles independently of cell division as well as to mid-cell when cells started constricting (Fig. 5A). As expected, neither polar nor mid-cell PilQ-sfGFP recruitment was observed in the absence of Tgl (Fig. 5A and Fig. S2A). Consistently, we also observed that upon induction of untagged PilQ synthesis in the Δ*pilQ* strain additionally expressing Tgl-sfGFP from the native site, Tgl-sfGFP transiently formed clusters at both poles (Fig. 5B and Fig. S2B). Finally, to determine whether PilQ recruited to the poles independently of cell division was competent to guide the assembly of the remaining components of the T4aPM, we repeated the PilQ induction experiment in a strain additionally expressing an active mCherry-PilM fusion (12) from the native site. The cytoplasmic PilM protein (Fig. 1) is the last component to be incorporated into the polar T4aPM in *M. xanthus* (27). Before induction of PilQ synthesis, mCherry-PilM localized diffusely to the cytoplasm; importantly, upon induction of *pilQ* expression, PilM also localized in a bipolar pattern (Fig. 5B and Fig. S2C).

**Figure 5.**
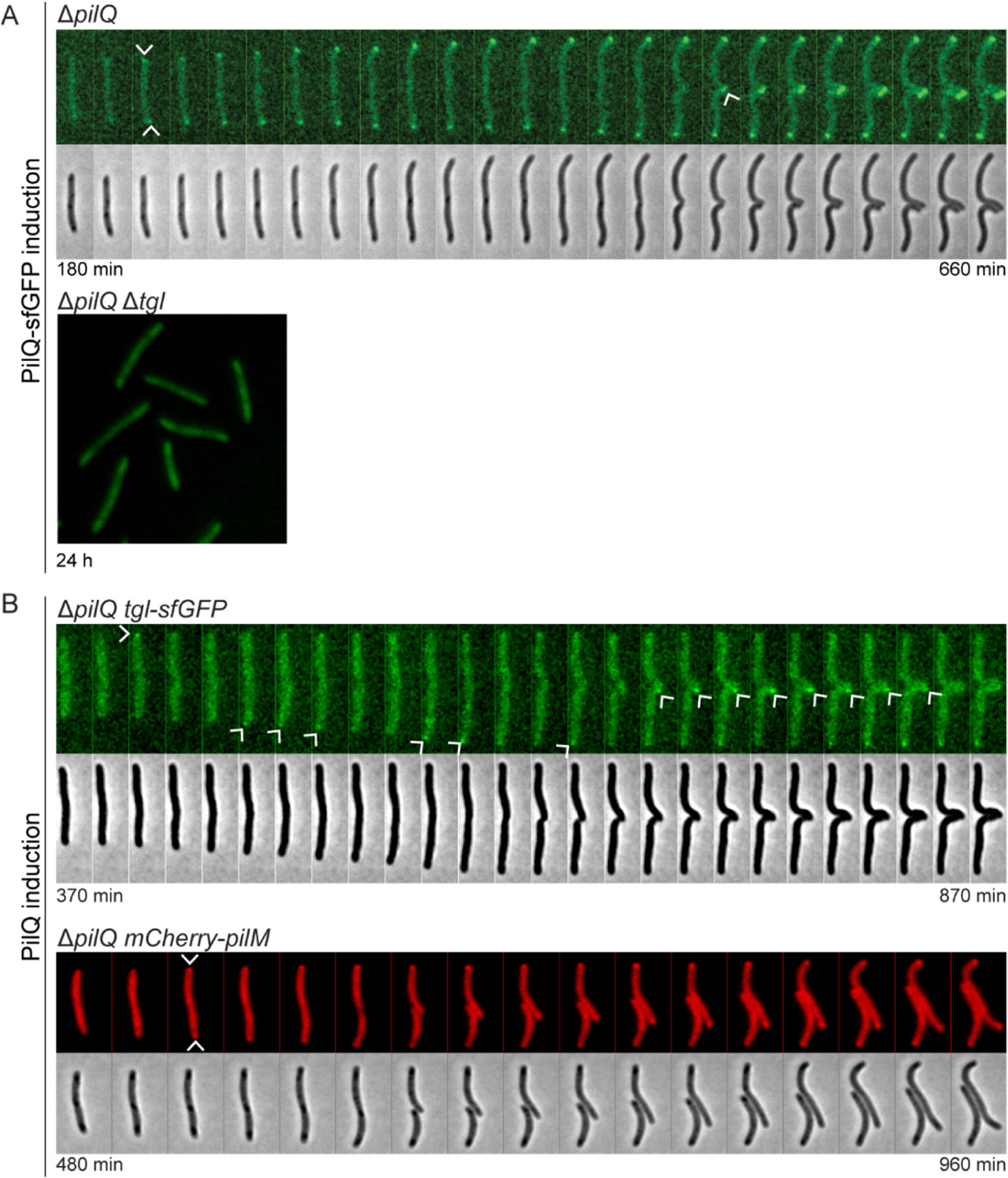
Cell division-independent assembly of T4aP machines (A, B) Induction of expression of *pilQ-sfGFP* (A) or *pilQ* (B) from P_van_ in the indicated strains followed by time-lapse epifluorescence microscopy. Time indicates interval after the addition of vanillate (t=0). In A, upper panel, the arrows indicate the first appearance of fluorescent clusters of PilQ-sfGFP. In A, lower panel, cells are shown after 24 hrs of PilQ-sfGFP induction in the Δ*tgl* mutant. In A, 10 µM vanillate was used for inducing PilQ-sfGFP accumulation at WT levels in the Δ*pilQ* background (Fig. S4A), and 500 µM vanillate was used to highly induce PilQ-sfGFP accumulation in the Δ*tgl* Δ*pilQ* background (Fig. S4A). In B, upper panel, 20 µM vanillate was used for inducing PilQ accumulation at WT levels in Δ*pilQ* cells expressing *tgl-sfGFP* (Fig. S4B). Arrows indicate time points at which the Tgl-sfGFP cluster is clearly visible. In B, lower panel 1mM vanillate was used to rapidly induce *pilQ* expression in cells co-expressing *mCherry-pilM* (Fig. S4C). Arrows indicate the first appearance of fluorescent clusters of mCherry-PilM. Localization of mCherry-PilM at the poles is used as a proxy to study correct assembly of the T4aPM. In A and B, strains analyzed contain the Δ*gltB* mutation.

Based on these observations, we conclude that the polar recruitment and OM incorporation of PilQ can occur at both poles independently of cell division, and that these secretins support the assembly of the remaining components of the T4aPM. Because this incorporation depends on Tgl, we also conclude that the cell division-independent PilQ incorporation into the OM follows the same mechanism as in the case of its incorporation at nascent and new poles.

### Tgl is not important for PilQ transport across the periplasm

To evaluate whether OM localization of Tgl is important for its function, we generated a strain expressing Tgl^C20G^-sfGFP (using the numbering of the full-length protein), in which the conserved Cys residue (+1 in the mature protein) (Fig. S3A) was substituted to Gly to prevent its acylation and, therefore, transport to and anchoring in the OM. Additionally, because an Asp in position +2 of mature lipoproteins in *Escherichia coli* can cause their retention in the IM (59), we also generated a strain expressing Tgl^S21D^-sfGFP. Expression of Tgl^C20G^-sfGFP and Tgl^S21D^-sfGFP from the native site or under the control of P_van_ only resulted in very low levels of accumulation of the proteins (Fig. S3B), thus precluding their further analyses.

Therefore, to obtain more insights into the function of Tgl, we determined the subcellular localization of PilQ in cell fractionation experiments in the presence and absence of Tgl. In wild-type (WT) cells, the PilQ monomer and multimer were enriched in the membrane fraction (Fig. 6A). Similarly, in Δ*tgl* cell extracts, monomeric PilQ was enriched in the membrane fraction. Control proteins previously shown to localize to the IM, OM or cytoplasm documented that the fractionation procedure worked properly (Fig. 6A).

**Figure 6.**
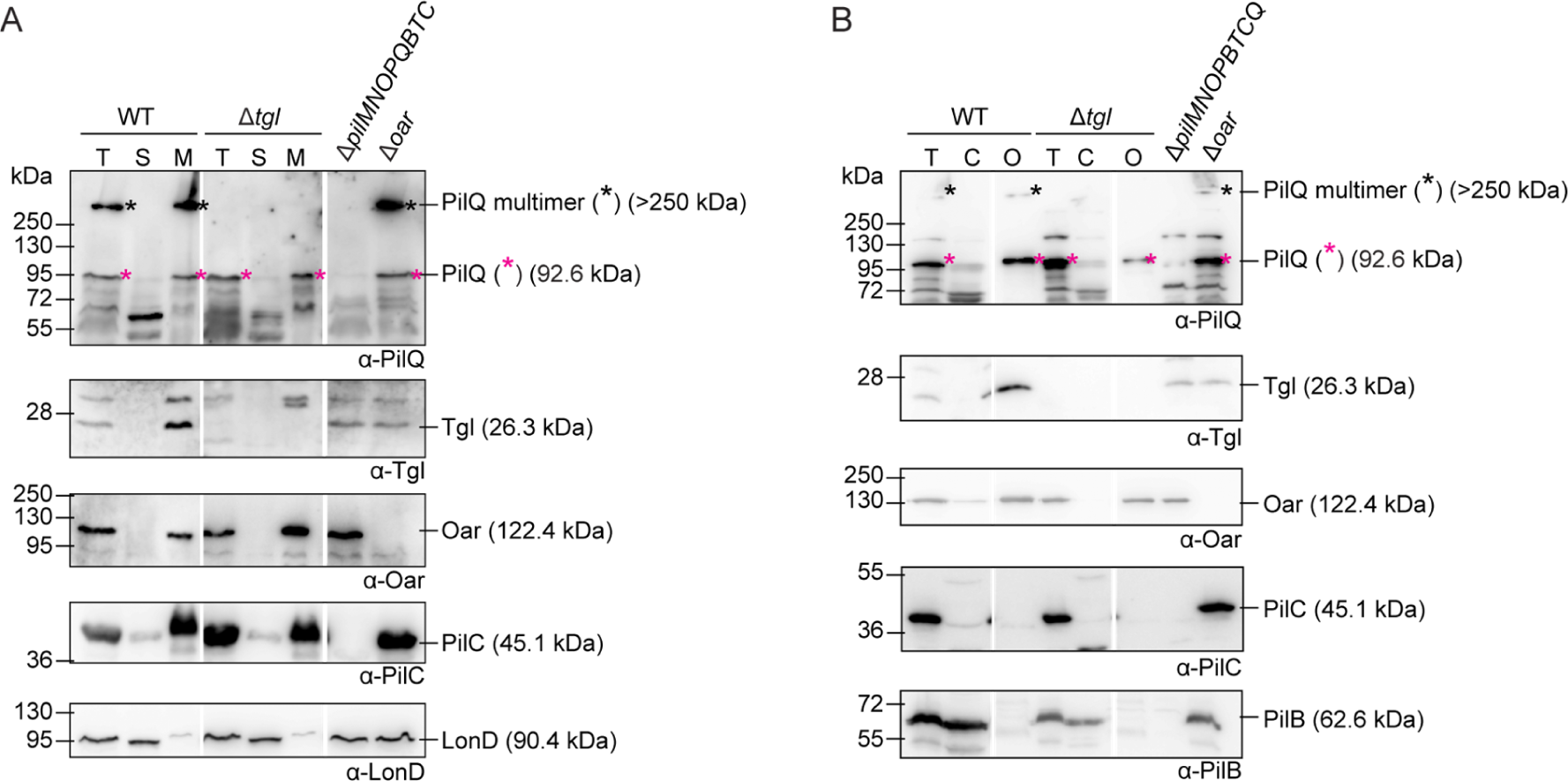
Subcellular localization of monomeric and multimeric PilQ as well as Tgl. (A) Total cell extracts (T) were fractionated into fractions enriched for soluble (S) and membrane (M) proteins. (B) Total cell extracts (T) were fractionated into fractions enriched for cytoplasmic (C), and OM (O) proteins. Protein from the same number of cells was loaded per lane and analyzed by immunoblotting. Oar is an OM protein (98), PilC is an IM protein (26), and LonD and PilB are cytoplasmic proteins (26, 99). These proteins served as controls that the fractionation procedure worked properly. Monomeric and oligomeric forms of PilQ are marked with an asterisk. Calculated molecular weights of proteins without signal peptide (if relevant) are indicated. Gaps indicate lanes removed for presentation purposes.

To determine whether monomeric PilQ is integrated in the IM or OM, we isolated the OM of WT and Δ*tgl* cells after osmotic shock with sucrose and EDTA treatment. Monomeric PilQ was detected in the OM fraction of both strains (Fig. 6B). As expected, in WT extracts, the heat- and detergent-resistant oligomers were also enriched in the OM fraction, while controls fractionated as expected (Fig. 6B). These results demonstrate that Tgl is not required for the transport of monomeric PilQ across the periplasm to the OM, and that monomeric PilQ is at to the OM.

### A computational structural model of the Tgl/PilQ complex

To evaluate how Tgl interacts with monomeric PilQ to promote its stability and multimerization in the OM, we analyzed the two proteins *in silico*. While the sequences of T4aPM pilotins are not well conserved (39) (Fig. S3A), it was previously suggested that they all share a similar superhelix structure composed of six TPR motifs (45, 60), which are typically involved in protein-protein interactions (61). In agreement with this suggestion, a high-confidence AlphaFold-based structure of monomeric Tgl includes 13 anti-parallel α-helices, among which helices 1 to 12 fold into six TPR motifs forming a superhelix (Fig. 7A and Fig. S3C). Additionally, the Tgl structural model could readily be superimposed on the solved structures of PilF and PilW (pilotins of the secretin of the T4aPM in *P. aeruginosa* and *N. meningitidis,* respectively (60, 62)) (Fig. 7B). While PilF does not contain disulfide bridges and PilW contains one, which is crucial for its function and connects TPR domains 3 and 4 (Fig. S3A) (60, 63), Tgl is predicted to contain three disulfide bridges that link TPR domains 5 and 6 as well as TPR 6 and α-helix 13 (Fig. 7A and Fig. S3A). Conservation analysis of the amino acid sequence of Tgl homologs using ConSurf revealed two conserved hydrophobic surfaces, one in the N-terminal TPR1 (from hereon CS1) and one within the concave groove of Tgl close to CS1 (from hereon CS2) (Fig. 7C).

**Figure 7.**
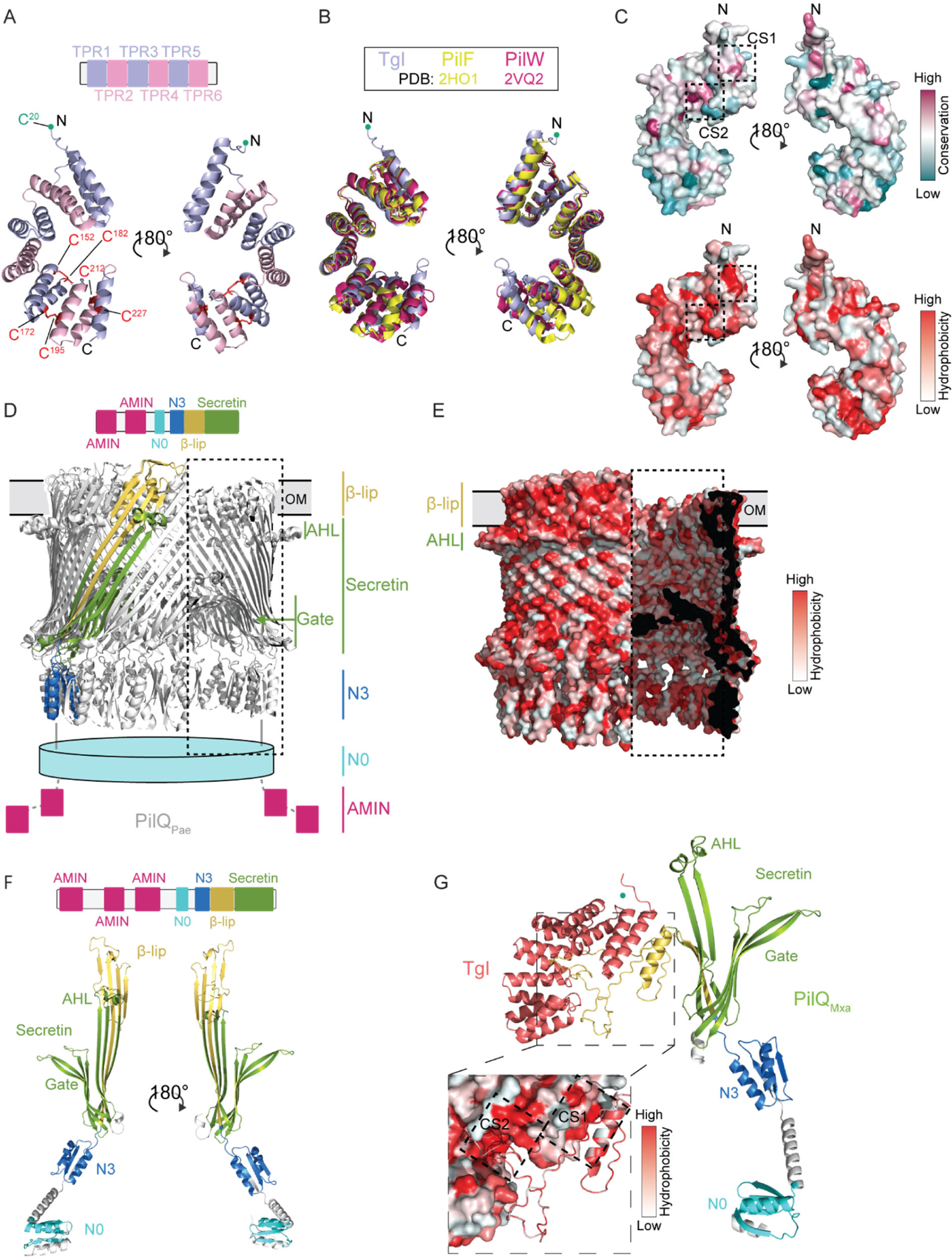
Structural characterization of Tgl alone and in complex with its PilQ secretin partner (A) AlphaFold model of mature Tgl. Upper panel, TPR domains 1 to 6 in Tgl are indicated to scale. Lower panel, AlphaFold structure of Tgl. Cys residues (Cys152/Cys182, Cys172/Cys195, Cys212/Cys227, numbering for unprocessed protein) engaged in disulfide bridge formation are indicated in red. Amino acids are indicated using the numbering of the full-length protein. (B) Superposition of the AlphaFold Tgl structure with the solved structures of PilF (PDB: 2HO1) (39) and PilW (PDB: 2VQ2) (60). (C) Surface representation of the sequence conservation calculated with ConSurf using 279 homolog sequences (upper panel) and hydrophobicity of Tgl (lower panel). The conserved hydrophobic surfaces CS1 in TPR1 and CS2 in the concave groove are marked with dashed squares. (D) CryoEM structure of the tetradecameric PilQ secretin of *P. aeruginosa* (PDB: 6VE2) (35). Upper panel, domain architecture of PilQ_Pae_. Lower panel, CryoEM structure, in which the different domains of one PilQ_Pae_ protomer are indicated as in (34) are colored as N3 (marine blue), secretin (green), and β-lip region (yellow). The N0 and two AMIN domains, which are not resolved in the structure, are represented by a cyan cylinder and magenta boxes, respectively. In the part of the secretin marked by the dashed box, the front part of the barrel structure has been removed to show the inside of the barrel with the gate.(E) Surface representation of the hydrophobicity of the cryoEM solved structure of the tetradecameric PilQ_Pae_ secretin. Note the amphipathic AHL and β-lip. In the part of the secretin marked by the dashed box, the front part of the barrel structure has been removed to show the inside of the barrel with the gate. Regions in which the protein structure was sliced are colored black. (F) AlphaFold model of *M. xanthus* PilQ monomer. Upper panel, domain architecture of PilQ. Lower panel, AlphaFold model with regions colored according to the domains and as described for panel D. For clarity, AMIN domains are not shown. (G) AlphaFold model of heterodimer of *M. xanthus* PilQ monomer and Tgl. Inset: Interaction interface between Tgl (surface representation) and PilQ (cartoon) colored according to hydrophobicity. The conserved hydrophobic surfaces CS1 in TPR1 and CS2 in the concave groove are marked with a dashed square. In A, B and G, the acylated N-terminal Cys residue of mature Tgl (residue Cys20 in the unprocessed protein) that places the protein at the inner leaflet of the OM is indicated by a green circle.

In secretins, the two or more conserved N-terminal N-domains are involved in oligomerization (64), interaction with IM components of the T4aPM (65), and also form part of the periplasmic vestibule, while the AMIN domains, if present, bind PG (24, 32, 34) (Fig. 7D). In the C-terminal region, the secretin domain (PF00263) forms β-sheets, which in the secretin oligomer form the gate, part of the periplasmic vestibule, and the amphipathic helical loop (AHL) (34, 37) (Fig. 7D). The hydrophobic surface of the AHL associates with the inner leaflet of the OM (Fig. 7D and E) (37). The amphipathic β-lip region in the C-terminal region forms part of the β-barrel (34), but mainly forms the β-stranded region with which the secretin spans the OM (Fig. 7D and E) (37).

Similar to other T4aPM secretins, PilQ from *M. xanthus* (PilQ_Mxa_) is divided into four main regions: the three AMIN domains connected by flexible linkers, the N0- and N3-domains, the β-lip region and the C-terminal secretin domain (Fig. 7F). Monomeric PilQ_Mxa_ was modeled with high confidence using AlphaFold (Fig. 7F and Fig. S3C) and could readily be superimposed on a protomer from the cryo-electron microscopy-based structure of *P. aeruginosa* PilQ (PilQ_Pae_) (35) (Fig. 7D and Fig. S4A). Similar results were obtained when using monomeric PilQ_Pae_ modeled with AlphaFold (Fig. S3C and Fig. S4A), supporting that the predicted structures are modeled with high confidence.

It is currently not known how pilotins of T4aPM secretins interact with their cognate secretin monomer. Therefore, to gain insights into how T4aPM secretins and their pilotins interact, we started with the *M. xanthus* proteins and used AlphaFold-Multimer to predict heterodimeric structures of Tgl and monomeric PilQ_Mxa_. Surprisingly, in the high-confidence heterodimer model, the amphipathic β-sheet of the β-lip observed in the structural model of the PilQ_Mxa_ monomer (Fig. 7F and Fig. S3C) is unfolded into (1) a hydrophobic α-helix and (2) an unstructured region (Fig. 7G). Remarkably, Tgl is modeled to specifically interact with this hydrophobic α-helix and this unstructured region via the conserved hydrophobic surfaces CS1 in TPR1 and CS2 in the concave groove, respectively (Fig. 7C and G). Underscoring the validity of this structural model of the heterodimer, PilF and PilW are also modeled with high confidence to associate with their partner secretin in the same way, i.e. using the same interfaces as in Tgl to contact the unfolded β-lip (Fig. S4B-D). Moreover, these specific interactions appear to depend on the cognate pilotin-secretin pair, because heterodimer modeling of Tgl with the *P. aeruginosa* or the *N. meningitidis* secretin yielded structural models of lower confidence, and in which some of these interactions were lost (Fig. S4E-G).

In conclusion, we suggest that T4aPM pilotins by associating with the unfolded β-lip of their cognate monomeric secretin keep this region, part of which will ultimately be inserted into the OM, in a conformation optimal for oligomerization and OM insertion. Once the secretin monomers multimerize and the correctly folded β-lip integrates into the OM, the interaction with the pilotin would be lost, thus explaining why the pilotin only transiently associates with the secretin.

### The presence and absence of AMIN domains in T4aPM secretins correlate with the piliation pattern

As shown here, septal and polar recruitment of the PilQ secretin, and therefore the T4aPM, in *M. xanthus* depends on its AMIN domains. Because AMIN domains are not universally conserved in T4aPM secretins (34), we wondered whether their presence or absence correlated with the localization pattern of the T4aP/T4aPM in other species. To this end, we selected bacteria with different T4aP localization patterns and studied the domain architecture of their secretin.

The PilQ secretins of *M. xanthus*, *P. aeruginosa*, *T. vulcanus* and *N. punctiforme* that localize to the cell poles (17, 18, 24) have three, two, one and one AMIN domains, respectively (Fig. 8A-D, Fig. S3C and Fig. S5). By contrast, the spherical cells of *Synechocystis* sp. PCC6803*, M. catarrhalis* as well as the rod-shaped cells of *B. cepacia* have PilQ homologs without AMIN domain (Fig. 8E-G and Fig. S5) and these species have peritrichous T4aP localization patterns (19, 22, 23). Interestingly, the coccoid *N. meningitidis* and *N. gonorrhoeae,* which assemble peritrichous T4aP (20, 21), both have PilQ homologs with two AMIN domains (Fig. 8H and Fig. S5). Finally, the PilQ homolog of *A. baylyi* contains two AMIN domains (Fig. 8I and Fig. S5), and has the unique lateral T4aP localization pattern (6).

**Figure 8.**
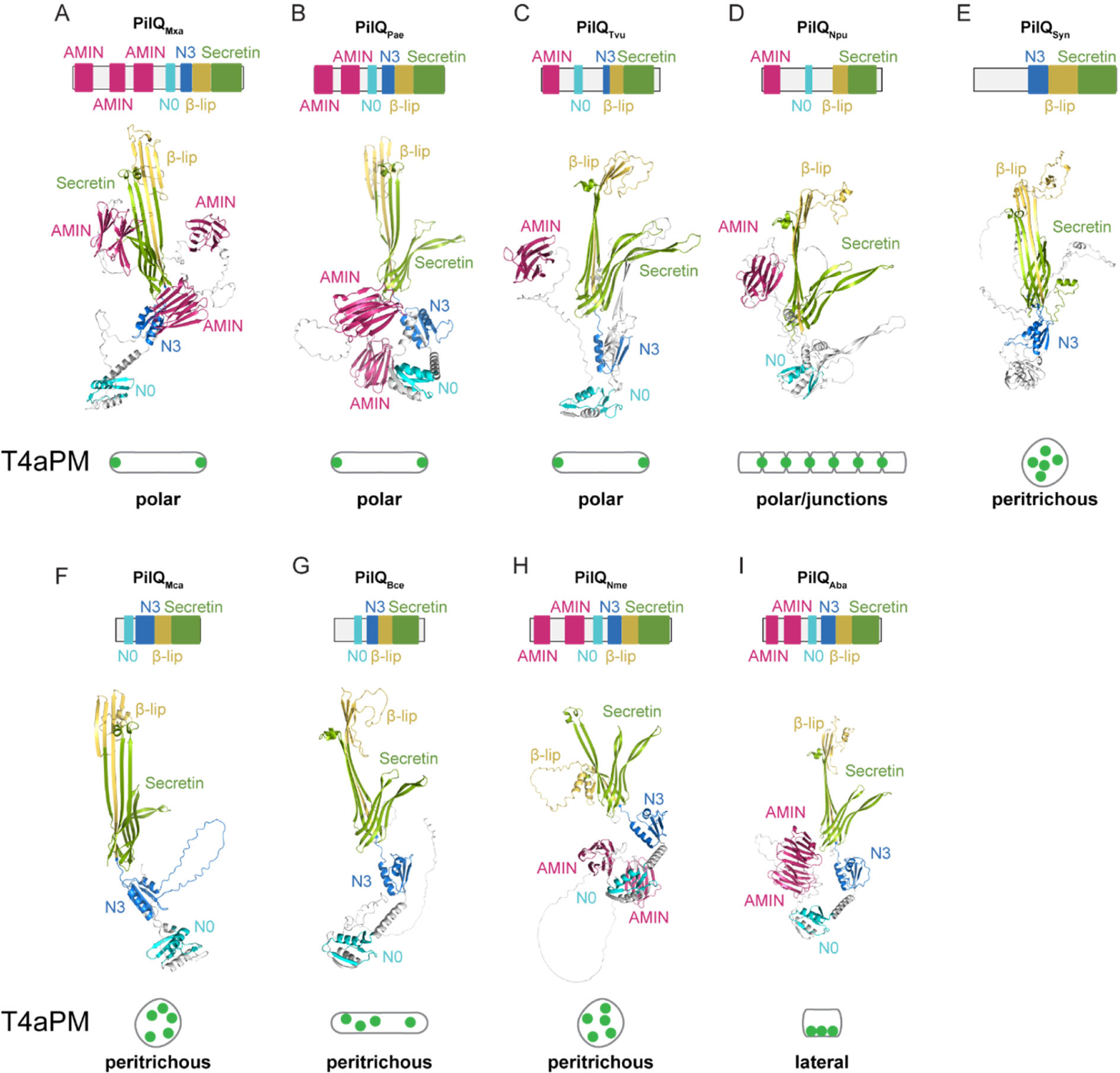
Characterization of PilQ secretins in other bacteria (A-I) Domain architecture and AlphaFold models of secretin monomers from (A) *M. xanthus* (PilQ_Mxa_), (B) *P. aeruginosa* (PilQ_Pae_) (GenBank: AAA16704.1), (C) *T. vulcanus* (PilQ_Tvu_) (GenBank: BAY52454.1), (D) *N. punctiforme* (PilQ_Npu_) (GenBank: RCJ37220.1), (E) *Synechocystis* sp. PCC6803 (PilQ_Syn_) (GenBank: BAA18278.1), (F) *M. catarrhalis* (PilQ_Mca_) (GenBank: ADG61696.1), (G) *B. cepacia* (PilQ_Bce_) (GenBank: ALK17307.1), (H) *N. meningitidis* (PilQ_Nme_) (GenBank: AHW75028.1), and (I) *A. baylyi* (PilQ_Aba_) (GenBank: AAK00351.1). The domains of PilQ are colored as described in Fig. 7D. T4aPM distribution within the cell is indicated (see text).

## Discussion

Here, we focused on the polar incorporation of the T4aPM in the rod-shaped cells of *M. xanthus* to understand how different localization patterns of T4aP are ultimately established. *M. xanthus* is an ideal system to address this question because the T4aPM assembly pathway is well-understood and initiates with the PilQ secretin in the OM (8, 27). Thus, using PilQ as a proxy for the T4aPM allowed us to examine how the specific localization of the T4aPM is determined. We demonstrate that PilQ is recruited to and begins to assemble in the OM at the nascent pole during cytokinesis, and these processes continue for 60-90 min in the two daughter cells after completion of cytokinesis. The recruitment and assembly eventually result in the symmetric localization of PilQ at the two cell poles. Consistent with the pilotin Tgl being important for PilQ multimer formation (27, 29), we observed that Tgl transiently associated with the nascent and new cell poles in a PilQ-dependent manner and largely in parallel with PilQ recruitment and OM assembly. Moreover, we demonstrate that PilQ recruitment to and assembly at the nascent and new poles depends on its PG-binding AMIN domains.

How, then, does PilQ assembled ato the nascent and new poles? Several lines of evidence support that this assembly is a two-step process that crucially depends on the PG-binding AMIN domains in PilQ in concert with Tgl. Firstly, a PilQ variant lacking all three AMIN domains accumulated but was not recruited to the nascent and new poles and did not assemble to form multimers, while a PilQ variant with only one AMIN domain is sufficient for correct assembly and polar localization of the T4aPM (8). In agreement with this observation, recruitment of the PilQ secretin in *P. aeruginosa* to the nascent poles was suggested to depend on its AMIN domains (24). AMIN domains also have crucial functions in the recruitment of the amidases AmiB and AmiC to the site of cell division in *E. coli* (56, 66) and the AmiC AMIN domain specifically binds to septal PG during cytokinesis (56–58). Secondly, previous work concluded that PG modifications acquired during cytokinesis are retained at the poles indefinitely (55). Thirdly, the pilotin Tgl, which is an OM lipoprotein, is important for PilQ stability and multimerization in the OM. Fourthly, in cells lacking PilQ, Tgl still accumulated; however, it did not localize to the nascent and new poles. Fifthly, the PilQ multimerization defect in Δ*tgl* cells can be extracellularly complemented by transfer of Tgl from Tgl^+^ cells (45, 48–50). Based on these lines of evidence, we suggest a model for the polar recruitment and OM incorporation of PilQ and, consequently, the complete T4aPM. In this model, PilQ monomers and Tgl are translocated to the OM independently of each other (Fig. 9, step 1). During and immediately after cytokinesis, PilQ monomers specifically recognize and stably bind to septal and polar PG via their AMIN domains (Fig. 9, step 2). These PilQ monomers either bring along Tgl or recruit Tgl to the poles (Fig. 9, step 2). The high local concentration of PilQ/Tgl complexes at the nascent and new poles, facilitates Tgl-dependent OM incorporation and multimerization of PilQ eventually resulting in the release of Tgl (Fig. 9, step 3-4). Upon assembly of the PilQ multimer in the OM, the remaining components of the T4aPM are incorporated (Fig. 9, step 5). In agreement with our model, PilQ can be recruited to both poles and assemble multimers in a cell-division independent manner supporting that both cell poles have the properties required for recruitment and incorporation of the PilQ secretin in the OM independently of a cell division event. We note that not all PilQ is localized to the cell poles. We, therefore, suggest that PilQ monomer recruitment to the nascent and new poles represents an example of a diffusion-and-capture mechanism for polar protein localization (67, 68). In this mechanism, OM-associated PilQ monomers and Tgl diffuse in two dimensions until the PilQ monomers recognize and bind septal and polar PG that serve as a polar landmark, thereby enabling polar Tgl localization (Fig. 9, step 1-2). We speculate that the number of assembled PilQ multimers in the OM at the cell poles, and therefore the number of T4aPM at the cell poles, is limited by the availability of the specific septal and polar PG recognized by the PilQ AMIN domains. In the future, it will be important to determine the exact PG recognized by the PilQ AMIN domains. Similarly, it will be important to determine whether monomeric PilQ and Tgl are recruited as a complex or sequentially. The assembled T4aPM in *M. xanthus* has a width of 15-20 nm (8, 12) while the average pore size of PG has been estimated to ∼2 nm (69). While the incorporation of T4aPM in parallel with cytokinesis is compatible these different dimensions, it is more difficult to understand how the T4aPM would be incorporated after completion of cytokinesis and independently of cell division. Thus, it will also be important to address how the T4aPM is assembled post-divisionally.

**Figure 9.**
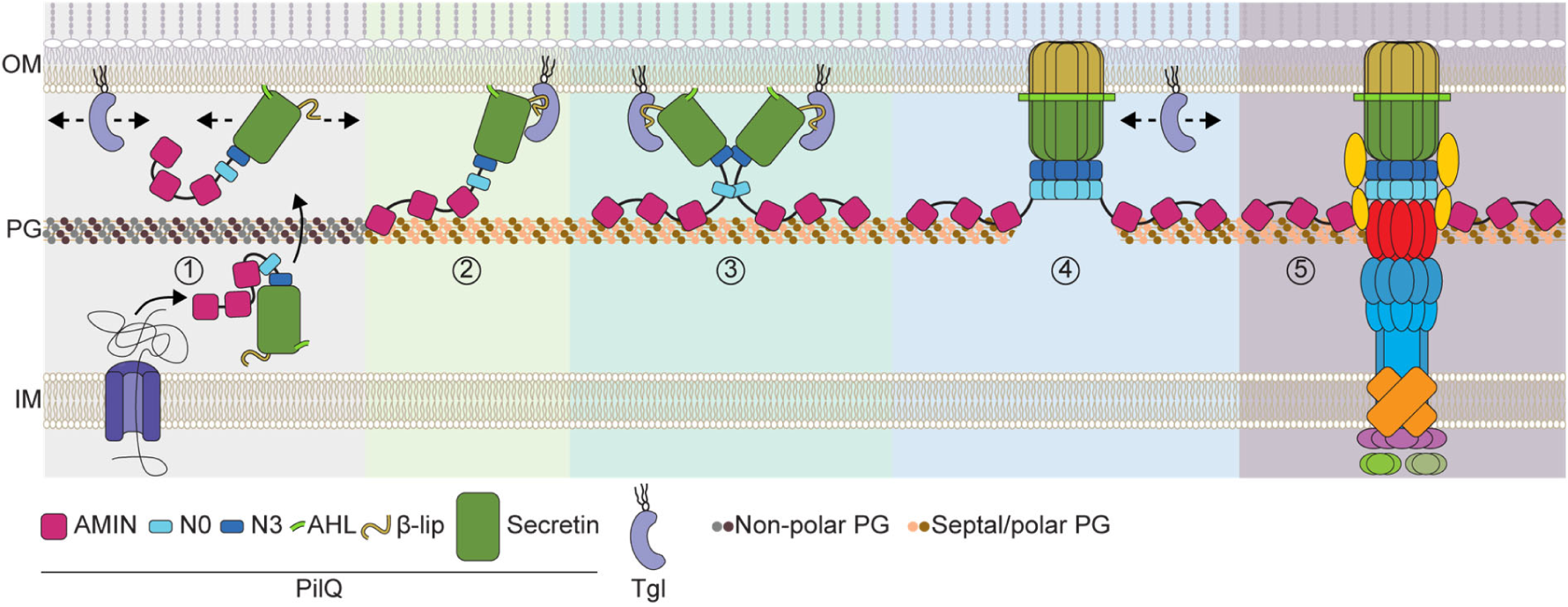
Model of polar incorporation and OM assembly of the PilQ secretin as well as polar assembly of the T4aPM in *M. xanthus* See main text for details. Note that in step 1, PilQ monomers and Tgl away from the septal and polar PG are shown not to interact; however, it is possible that the two proteins interact prior to their polar localization. In step 1 and 4, the arrows indicate that the proteins can diffuse in the OM. In step 1-3, PilQ is shown to associate with the OM via its AHL domain; however, it is not known how PilQ monomers associate with/integrate into the OM. In step 3, while the secretin oligomerizes from 12-15 PilQ monomers, only two are shown for illustration purposes. In step 5, all T4aPM components except for PilQ are colored as in Fig. 1.

How then does Tgl stimulate PilQ multimer formation in the OM? Tgl is important for PilQ stability and multimerization in the OM. In cellular fraction experiments, we found that the PilQ monomer is associated with the OM in a Tgl-independent manner demonstrating that Tgl is not required for translocating monomeric PilQ from the IM across the periplasm to the OM. We note that whether PilQ is associated with the OM or integrated into the OM cannot be distinguished based on these experiments. High confidence *in silico* structural models of monomeric Tgl, monomeric PilQ, and heterodimeric Tgl/PilQ complexes, support that Tgl interacts with monomeric PilQ via hydrophobic interfaces. Specifically, our structural models suggest that two conserved hydrophobic surfaces, i.e. CS1 in TPR1 and CS2 in the concave groove, in Tgl interact with the hydrophobic parts of the unfolded amphipathic β-lip of monomeric PilQ. Therefore, our results suggest that Tgl at the OM binds the OM associated PilQ monomer thereby (1) stimulating multimerization by maintaining an oligomerization-ready conformation of the PilQ monomer, (2) protecting monomeric PilQ from proteolytic degradation, and (3) ensuring that the assembled secretin only forms at the OM. Because Tgl is associated with the OM via its acylated N-terminus, CS1 and CS2 are close to the OM and, therefore, ideally positioned to assist in PilQ secretin integration into the OM. Once PilQ monomers multimerize and integrate into the OM, the interaction with Tgl would be lost because the β-lip is integrated into the OM and the interaction surfaces no longer available for interaction with Tgl (Fig. 9, step 3-4). *In silico* structural models of PilF/PilQ and PilW/PilQ heterodimers support that they form complexes similar to that of Tgl/PilQ. These models of the heterodimers are also supported by the observations that PilF interacts with the C-terminal region of monomeric PilQ_Pae_ and that loss-of-function PilF variants have substitutions in TPR1 (70). We, therefore, propose that T4aPM pilotins and their cognate monomeric secretin use the same conserved mechanism in which the pilotin interacts with the unfolded β-lip of the monomeric secretin to aid its OM integration. Lending further support for this generalized mechanism of the cognate T4aPM pilotin/secretin pairs, cognate pilotin/secretin pairs of the T2SS and T3SS interact via the so-called S-domain at the C-terminus of the secretin monomer (32, 33). However, T4aPM pilotins are structurally different from T2SS and T3SS pilotins (71), and T4aPM secretins lack the S-domain (32, 33).

Because different species have different T4aP patterns and the PilQ AMIN domains in polarly piliated *M. xanthus* (here) and *P. aeruginosa* (24) are essential for the polar assembly of the T4aPM, we asked whether there is a correlation between piliation patterns and the presence/absence of AMIN domains in the relevant secretins. Interestingly, we found that not only the PilQ secretins of *M. xanthus* and *P. aeruginosa* but also of the polarly piliated *T. vulcanus* and *N. punctiforme* contain AMIN domains, while the secretins of the peritrichiously piliated *Synechocystis* sp. PCC6803*, M. catarrhalis* and *B. cepacia* cells do not contain AMIN domains. In this survey, three species did not follow the overall correlation between piliation pattern and the presence/absence of AMIN domain(s) in the relevant secretin. Specifically, *N. meningitidis* and *N. gonorrhoeae* assemble peritrichous T4aP and both have secretins that contain AMIN domains. Notably, it has been suggested that these two species have a rod-shaped ancestor and that the emergence of their coccoid cell shape derives from relatively recent gene losses from the genome of this ancestor (72, 73). Thus, we speculate that the AMIN domains in the *N. meningitidis* and *N. gonorrhoeae* T4aPM secretins are remnants from the rod-shaped ancestor. The third species in which the piliation pattern and absence /presence of AMIN domain(s) correlation did not match was *A. baylyi.* This species has a unique lateral piliation pattern and its T4aPM secretin contains two AMIN domains, suggesting that *A. baylyi* may potentially accumulate a specific form of PG laterally that is recognized by its T4aPM secretin. Interestingly, the secretins of T2SS and T3SS lack AMIN domains and have been reported to have a dispersed localization (74–76). Thus, the presence/absence of AMIN domains in the relevant secretin for the localization of the relevant macromolecular structure may extend beyond the T4aPM.

## Supporting information

All Supplementary Information

## Acknowledgment

The authors thank Martin Thanbichler for helpful discussions, Luís Antonío Menezes Carreira for constructing pLC220 and Anke Treuner-Lange for generating strain SA11377. This work was generously supported by the Max Planck Society (to LS-A).

## Availability of data and materials

The authors declare that all data supporting this study are available within the article and its Supplementary Information files.

## Materials and Methods

### Bacterial strains and growth media

All *M. xanthus* strains are derivatives of the wild type DK1622 (15) and are listed in Table 1. In-frame deletions and gene replacements were generated as described (77) and were verified by PCR. Point mutation replacements were confirmed by DNA sequencing. *M. xanthus* cells were grown at 32°C in 1% CTT broth (1% (w/v) Bacto Casitone, 10 mM Tris-HCl pH 8.0, 1 mM K_2_HPO_4_/KH_2_PO_4_ pH 7.6, and 8 mM MgSO_4_) or on 1% CTT 1.5% agar (50) supplemented when required with kanamycin (50 µg ml^−1^) or oxytetracycline (10 µg ml^−1^).

**Table 1.** *M. xanthus* strains used in this work

| Strain | Genotype | Reference |
| --- | --- | --- |
| DK1622 | WT | (15) |
| DK10410 | $\Delta pilA$ | (100) |
| DK8615 | $\Delta pilQ$ | (101) |
| DK10405 | $\Delta tgl::tet^R$ | (46, 49) |
| SA6053 | $\Delta tgl$ | (27) |
| SA6024 | $\Delta pilBTCMNOPQ$ | (27) |
| SA3922 | $\Delta gltB$ | (85) |
| SA7192 | $pilQ::pilQ-sfGFP$ | (5) |
| SA11377 | $\Delta oar$ | This study |
| SA12016 | $tgl::tgl-sfGFP$ | This study |
| SA12017 | $pilQ::pilQ-sfGFP \Delta gltB$ | This study |
| SA12021 | $tgl::tgl-sfGFP \Delta gltB$ | This study |
| SA12031 | $\Delta tgl::tet^R pilQ::pilQ-sfGFP$ | This study |
| SA12032 | $\Delta pilQ tgl::tgl-sfGFP$ | This study |
| SA12035 | $tgl::tgI^{C20G}-sfGFP$ | This study |
| SA12047 | $\Delta tgl 18-19::P_{van} tgl-sfGFP$ | This study |
| SA12048 | $\Delta tgl 18-19::P_{van} tgI^{C20G}-GFP$ | This study |
| SA12049 | $pilQ::pilQ^{\Delta AMIN \times 3 (\Delta 31-475)}-sfGFP$ | This study |
| SA12050 | $\Delta tgl pilQ::pilQ-sfGFP$ | This study |
| SA12054 | $\Delta pilQ \Delta gltB 18-19::P_{van} pilQ-sfGFP$ | This study |
| SA12073 | $\Delta tgl 18-19::P_{van} tgI^{S21D}-sfGFP$ | This study |
| SA12074 | $pilQ::pilQ^{AMIN \times 3 (1-475)}-sfGFP$ | This study |
| SA12078 | $\Delta pilQ tgl::tgl-sfGFP \Delta gltB 18-19::P_{van} pilQ$ | This study |
| SA12085 | $\Delta gltB \Delta tgl \Delta pilQ 18-19::P_{van} pilQ-sfGFP$ | This study |
| SA12088 | $\Delta pilQ \Delta gltB pilM::mCherry-pilM 18-19::P_{van} pilQ$ | This study |

Plasmids used in this study are listed in Table 2. Plasmids were propagated in *E. coli* Mach1 (Δ*recA*1398 *endA*1 *tonA* Φ80Δ*lacM*15 Δ*lacX*74 *hsdR* (r_K_^−^ m_K_^+^); Invitrogen), which was grown at 37°C in lysogeny broth (10 mg tryptone ml^−1^, 5 mg yeast extract ml^−1^ and 10 mg NaCl ml^−1^) supplemented when required with kanamycin (50 µg ml^−1^).

**Table 2.** Plasmids used in this work

| Plasmid | Description | Reference |
| --- | --- | --- |
| pBJ114 | Km<sup>r</sup> *galK* | (102) |
| pMR3690 | Km<sup>r</sup>, P<sub>van</sub> | (103) |
| pDK25 | pBJ114, for generation of a *gltB* in-frame deletion, Km<sup>r</sup> | (85) |
| pNG020 | pBJ114, for generation of an *oar* in-frame deletion, Km<sup>r</sup> | (98) |
| pAP37 | pBJ114, for native site replacement of *pilQ* with *pilQ-sfGFP*, Km<sup>r</sup> | (5) |
| pMAT123 | pBJ114, for generation of a *pilQ* in-frame deletion, Km<sup>r</sup> | (104) |
| pMAT336 | pBJ114, for native site replacement of *pilM* with *mCherry-pilM*, Km<sup>r</sup> | (12) |
| pLC220 | pBJ114, for native site replacement of *tgl* with *tgl-sfGFP*, Km<sup>r</sup> | This study |
| pMH111 | pBJ114, for native site replacement of *tgl* with *tgl<sup>C20G</sup>*, Km<sup>r</sup> | This study |
| pMH118 | pMR3690, induction construct of *pilQ-sfGFP* expressed from the vanillate promoter, Km<sup>R</sup> | This study |
| pMH119 | pMR3690, induction construct of *tgl-sfGFP* expressed from the vanillate promoter, Km<sup>R</sup> | This study |
| pMH120 | pMR3690, induction construct of *tgl<sup>C20G</sup>-sfGFP* expressed from the vanillate promoter, Km<sup>R</sup> | This study |
| pMH121 | pBJ114, for generation of an in-frame deletion of the three AMIN domains of *pilQ-sfGFP* (Δ31-475), Km<sup>r</sup> | This study |
| pMH122 | pBJ114, for generation of a *tgl* in-frame deletion, Km<sup>r</sup> | This study |
| pMH125 | pBJ114, for native site replacement of *pilQ-sfGFP* with *pilQ<sup>AMIN×3</sup>* (1-475)-sfGFP, Km<sup>r</sup> | This study |
| pMH127 | pMR3690, induction construct of *tgl<sup>S21D</sup>-sfGFP* expressed from the vanillate promoter, Km<sup>R</sup> | This study |
| pMP183 | pMR3690, induction construct of *pilQ* expressed from the vanillate promoter, Km<sup>R</sup> | This study |

### Plasmid construction

All oligonucleotides used are listed in Table S1. All constructed plasmids were verified by DNA sequencing.

For **pLC220** (plasmid for replacement of *tgl* with *tgl-sfGFP* in the native site): the *tgl-sfGFP* fragment was amplified from pSC104 (27) using primers tgl_fw_hindiii/sfgfp_rv_xbaI. The downstream fragment was amplified from genomic DNA from *M. xanthus* DK1622 using the primer pair tgl_ds_fw_xbaI/ tgl_ds_rv. To generate the full-length insert, both DNA fragments were digested with XbaI and ligated. Next, the insert was digested with HindIII and EcoRI, and cloned into pBJ114.

For **pMH111** (plasmid for replacement of *tgl* with *tgl*^C20G^ at the native site): up- and downstream fragments were amplified using genomic DNA from *M. xanthus* DK1622 as DNA template and the primer pairs tgl_CtoG_A_HindIII/tgl_CtoG_Bov and tgl_CtoG_Cov/ tgl_CtoG_D_BamHI, respectively. To generate the full-length insert, an overlapping PCR using the two fragments as DNA templates and the primer pair tgl_CtoG_A_HindIII/ tgl_CtoG_D_BamHI was performed. Subsequently, the fragment was digested with HindIII and BamHI, and cloned into pBJ114.

For **pMH118** (plasmid for expression of *pilQ-sfGFP* from the *18-19* site under the control of the vanillate promoter): *pilQ-sfGFP* was amplified using genomic DNA from *M. xanthus* SA7192 (*pilQ::pilQ-sfGFP*) (5) as DNA template and the primer pair Pvan_PilQ_fwd_NdeI/ sfGFP_rev_pilQ_EcoRI. The fragment was digested with NdeI and EcoRI, and cloned into pMR3690.

For **pMH119** (plasmid for expression of *tgl-sfGFP* from the *18-19* site under the control of the vanillate promoter): *tgl-sfGFP* was amplified using genomic DNA from *M. xanthus* SA12016 (*tgl::tgl-sfGFP*) as DNA template and the primer pair Pvan_tgl_fw_NdeI/sfGFP_rv_tgl_EcoRI. The fragment was digested with NdeI and EcoRI, and cloned into pMR3690.

For **pMH120** (plasmid for expression of *tgl*^C20G^*-sfGFP* from the *18-19* site under the control of the vanillate promoter): *tgl*^C20G^*-sfGFP* was amplified using genomic DNA from *M. xanthus* SA12035 (*tgl*^C20G^*::tgl-sfGFP*) as DNA template and the primer pair Pvan_tgl_fw_NdeI/ sfGFP_rv_tgl_EcoRI. The fragment was digested with NdeI and EcoRI, and cloned into pMR3690.

For **pMH121** (for generation of an in-frame deletion of the AMIN×3 domains of native *pilQ*): up- and downstream fragments were amplified from genomic DNA from *M. xanthus* DK1622 using the primer pairs PilQ_dAMIN_A_XbaI/ PilQ_dAMIN_B and PilQ_dAMIN_C/ pilQ_dAMIN_D_HindIII, respectively. Subsequently, the up- and downstream fragments were used as a template for an overlapping PCR with the primer pair PilQ_dAMIN_A_XbaI/ pilQ_dAMIN_D_HindIII, to generate the AD fragment. The AD fragment was digested with XbaI and HindIII, and cloned in pBJ114.

For **pMH122** (for generation of an in-frame deletion of *tgl*): up- and downstream fragments were amplified from genomic DNA of SA6053 (Δ*tgl*) (27) using the primer pair tgl-A_XbaI/tgl-D_EcoRI to generate the AD fragment as described in (77). The AD fragment was digested with XbaI/EcoRI and cloned in pBJ114.

For **pMH125** (for replacement of *pilQ* with *pilQ*^AMINs×3^ (^1–475^)-*sfGFP* in the native site of the *pilQ::pilQ-sfGFP* strain): up- and downstream fragments were amplified from pMH118 using the primer pairs PilQAMIN_A_KpnI/ PilQAMIN_sfGFP_overlay_rev and PilQamin_sfGFP_overlay_fwd/ sfGFP_rev_pilQ_EcoRI, respectively. Subsequently, the up- and downstream fragments were used as a template for an overlapping PCR with the primer pair PilQAMIN_A_KpnI/ sfGFP_rev_pilQ_EcoRI, to generate the AD fragment. The AD fragment was digested with KpnI and EcoRI, and cloned in pBJ114.

For **pMH127** (plasmid for expression of *tgl*^S21D^*-sfGFP* from the *18-19* site under the control of the vanillate promoter): *tgl*^S21D^*-sfGFP* was amplified using pMH119 as DNA template and the primer pairs Pvan forw/Tgl_S21G_overlay_rev and Tgl_S21G_overlay_fwd/sfGFP_rv_tgl_EcoRI to introduce the point mutation. Subsequently, both PCR fragments were used as a template for an overlapping PCR with the primer pair Pvan forw/sfGFP_rv_tgl_EcoRI, to generate the full-length fragment. The fragment was digested with NdeI and EcoRI, and cloned into pMR3690.

For **pMP183** (plasmid for expression of *pilQ* from the *18-19* site under the control of the vanillate promoter): *pilQ* was amplified using pMH118 as DNA template and the primer pair Pvan_PilQ_fwd_NdeI/ PilQ_rev_EcoRI. The fragment was digested with NdeI and EcoRI, and cloned into pMR3690.

### Motility assays

T4aP-dependent motility assays were performed as described (78). Briefly, exponentially growing *M. xanthus* cultures were harvested (6,000 *g*, 3 min, RT) and resuspended in 1% CTT to a calculated density of 7×10^9^ cells ml^−1^. 5 µl aliquots were spotted on 0.5% CTT 0.5% select-agar (Invitrogen). After 24 h incubation at 32°C, cells were imaged using an M205FA Stereomicroscope (Leica) equipped with a Hamamatsu ORCA-flash V2 Digital CMOS camera (Hamamatsu Photonics), and images were analyzed using Metamorph® v 7.5 (Molecular Devices).

### Epifluorescence microscopy

Cells were visualized following a slightly modified protocol (79). Briefly, exponentially growing cells were placed on a glass coverslip attached to a metal frame. Cells were covered with a thick 1% agarose pad supplemented with 0.2% (w/v) Bacto Casitone and TPM (10 mM Tris-HCl pH 8.0, 1 mM K_2_HPO_4_/KH_2_PO_4_ pH 7.6, 8 mM MgSO4), and supplemented with vanillate or cephalexin as indicated. For long time-lapse microscopy, the pad was additionally sealed with parafilm to reduce evaporation. Cells were imaged using a DMi8 inverted microscope and a Hamamatsu ORCA-Flash4.0 V2 Digital CMOS C11440 or a DFC9000 GT (Leica) camera. Images were analyzed using Metamorph® v 7.5 (Molecular Devices) and ImageJ (Schindelin et al., 2012). Image segmentation was done using Omnipose (80) and cell outlines were transformed to Oufti-compatible meshes using Matlab R2020a (The MathWorks). Segmentation was manually curated using Oufti (81). For signal detection and background correction, a previously published Matlab script was used (82). Because the Tgl-sfGFP fluorescent clusters have low fluorescence intensity, the script was modified to detect the strongest pixel intensity in each cell segment assigned by Oufti. Specifically, each pixel intensity in each segment was normalized to the maximum pixel intensity within the cell. Next, to identify cells with one or more fluorescent cluster(s), cells were only considered to have a cluster if less than 10% of the selected pixel intensities had a normalized fluorescence above 0.75. Hence identifying cells with an intense and narrow fluorescent peak.

### Immunoblot analysis

Immunoblots were carried out as described (83). Rabbit polyclonal α-Tgl (dilution: 1:2,000) (27), α-PilQ (dilution, 1:5,000) (26), α-PilB (dilution: 1:2,000) (84), α-PilC (dilution: 1:2000) (26), α-Oar (1:10,000) (85) and α-LonD (dilution: 1:5,000) (12), were used together with a horseradish peroxidase-conjugated goat anti-rabbit immunoglobulin G (1:15,000) (Sigma) as a secondary antibody. Mouse α-GFP antibodies (dilution: 1:2,000) (Roche) were used together with horseradish peroxidase-conjugated sheep α-mouse immunoglobulin G (dilution: 1:2,000) (GE Healthcare) as a secondary antibody. Blots were developed using Luminata Forte Western HRP Substrate (Millipore) on a LAS-4000 imager (Fujifilm).

### Fractionation of *M. xanthus* cells

To fractionate *M. xanthus* cells into fractions enriched for soluble or membrane proteins, 20 ml of an exponentially growing *M. xanthus* suspension culture were harvested by centrifugation (8,000 *g*, 10 min, RT) and concentrated to an optical density at 550 nm (OD_550_) of 28 in resuspension buffer (50 mM Tris-HCl pH 7.6, 250 mM NaCl supplemented with Complete EDTA-free protease inhibitor (Roche)). Cells were lysed by sonication with 5×30 pulses, pulse 60%, amplitude 60% with a UP200St sonifier and microtip (Hielscher), and the lysate was cleared by centrifugation (12,000 *g,* 5 min, RT). As a sample for total cellular protein, an aliquot of the cleared lysate was taken and mixed with 4×SDS lysis buffer (200 mM Tris-HCl pH 6.8, 8% SDS, 40% glycerol, 400 mM DTT, 6 mM EDTA, 0.4% bromphenol blue). A 200 µl aliquot of the remaining supernatant was subjected to ultracentrifugation using an Air-Fuge (Beckman) (100,000 *g*, 20 min, RT). The resulting supernatant is enriched in soluble proteins and a sample was taken and mixed with 4×SDS lysis buffer. The pellet was washed by resuspension in 200 µl resuspension buffer and was subjected to ultracentrifugation as above. The remaining pellet, which is enriched in IM and OM membrane proteins, was resuspended in 100 µl 1×SDS lysis buffer. All samples were heated for 10 min at 95°C, separated by SDS-PAGE and analyzed by immunoblot.

### OM protein enrichment

As a sample for total cellular protein (total fraction), 2 ml of an exponentially growing *M. xanthus* cell suspension were harvested by centrifugation (8,000 *g*, 3 min, RT) and concentrated to an OD_550_ of 7 in 1× SDS lysis buffer. To isolate a fraction enriched for OM proteins, 50 ml of the cell suspension were harvested (4,700 *g*, 25 min, 4°C), and the pellet was gently resuspended in TSE8-buffer (100 mM Tris-HCl pH 8, 1 mM EDTA, 20% (w/v) sucrose, protease inhibitor cocktail (Roche)) to a concentration corresponding to OD_550_=50. The sample was incubated for 30 min at 4°C to release the OM, followed by centrifugation of the samples (16,000 *g*, 30 min, 4°C). The resulting supernatant is enriched in OM and periplasmic proteins and was recovered for the following steps, while the pellet, containing cells without OM or where the OM had not been released, was frozen at −20°C. Next, 150 µl of the supernatant was ultra-centrifuged using an Air-Fuge (Beckman) (∼133,000 *g*, 1 h, RT) to separate the OM from periplasmic proteins. The resulting supernatant was discarded and the OM-enriched pellet (OM fraction) was resuspended in 150 µl 1×SDS lysis buffer. The frozen pellet was thawed, resuspended to OD=50 in resuspension buffer (50 mM Tris pH 7.6, 10 mM MglCl_2_) and lysed by sonication. Cell debris was removed by centrifugation (16,000 *g*, 15 min, 4°C). The cell-free supernatant (∼150 µl) was subjected to ultra-centrifugation as described above. The resulting supernatant contained cytoplasmic proteins and was mixed with 4×SDS lysis buffer (cytoplasmic fraction). All samples were boiled 10 min at 95°C, separated by SDS-PAGE, and analyzed by immunoblot.

### Bioinformatics

Full-length protein sequences or sequences in which the signal peptide was identified with SignalP 6.0 (86) and removed, were used for AlphaFold and AlphaFold-Multimer modeling via ColabFold (1.3.0) (87–89). The predicted Local Distance Difference Test (pLDDT) and predicted Alignment Error (pAE) graphs of the five models generated were made using a custom Matlab R2020a (The MathWorks) script (90). Ranking of the models was performed based on combined pLDDT and pAE values, with the best-ranked models used for further analysis and presentation. Per residue model accuracy was estimated based on pLDDT values (>90, high accuracy; 70-90, generally good accuracy; 50-70, low accuracy; <50, should not be interpreted) (87). Relative domain positions were validated by pAE. The pAE graphs indicate the expected position error at residue X if the predicted and true structures were aligned on residue Y; the lower the pAE value, the higher the accuracy of the relative position of residue pairs and, consequently, the relative position of domains/subunits/proteins (87). PyMOL version 2.4.1 (http://www.pymol.org/pymol) was used to analyze and visualize the models. Structural alignments were performed using the PyMOL Alignment plugin with default settings. Hydrophobicity was calculated in PyMol according to the hydrophobicity scale (91). Conservation of Tgl residues was assessed using the ConSurf server with default settings (92). Protein domains were identified using the Interpro server (93) and the predicted AlphaFold structures. The alignment of Tgl, PilF, and PilW was generated using Muscle5 (5.1) (94).

