## Supplementary material for "The mechanism for polar localization of the type IVa pilus machine": All Supplementary Information

**This file contains:**

- Supplementary Figures 1-5
- Table S1

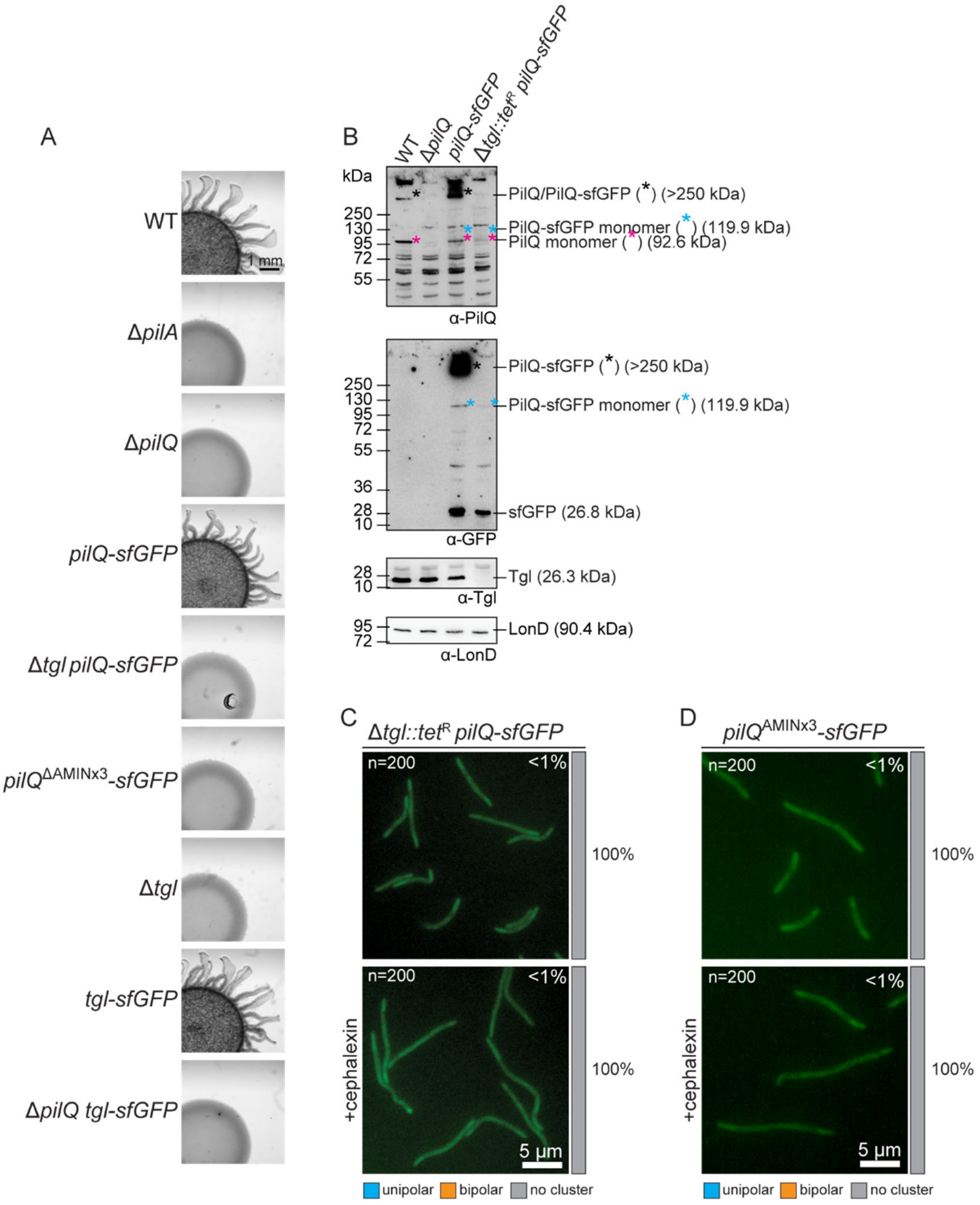


**Figure S1. Analysis of PilQ-sfGFP and Tgl-sfGFP variants functionality and localization**

(A) Colony-based motility assay of the indicated strains. T4aP-dependent motility was analyzed on 0.5% agar and images were recorded after 24 h. The Δ*pilA* mutant is deficient in T4aP-dependent motility and was used as the negative control. Scale bar: 1 mm.

(B) Immunoblot detection of PilQ-sfGFP in the Δ*tgl::Tet*$^{R}$ background strain. Protein from the same number of cells from exponentially growing suspension cultures was loaded per lane. Blot was probed with the indicated antibodies. The blot was stripped before applying a new antibody. LonD served as a loading control. Monomeric and oligomeric forms of PilQ/PilQ-sfGFP are marked with an asterisk. Calculated molecular weights of proteins without signal peptide (if relevant) are indicated.

(C-D) Localization of PilQ-sfGFP in the Δ*tgl::Tet*$^{R}$ background strain or of PilQ$^{AMINx3}$-sfGFP, and in the presence and absence of cephalexin as in Fig. 2B left panel.

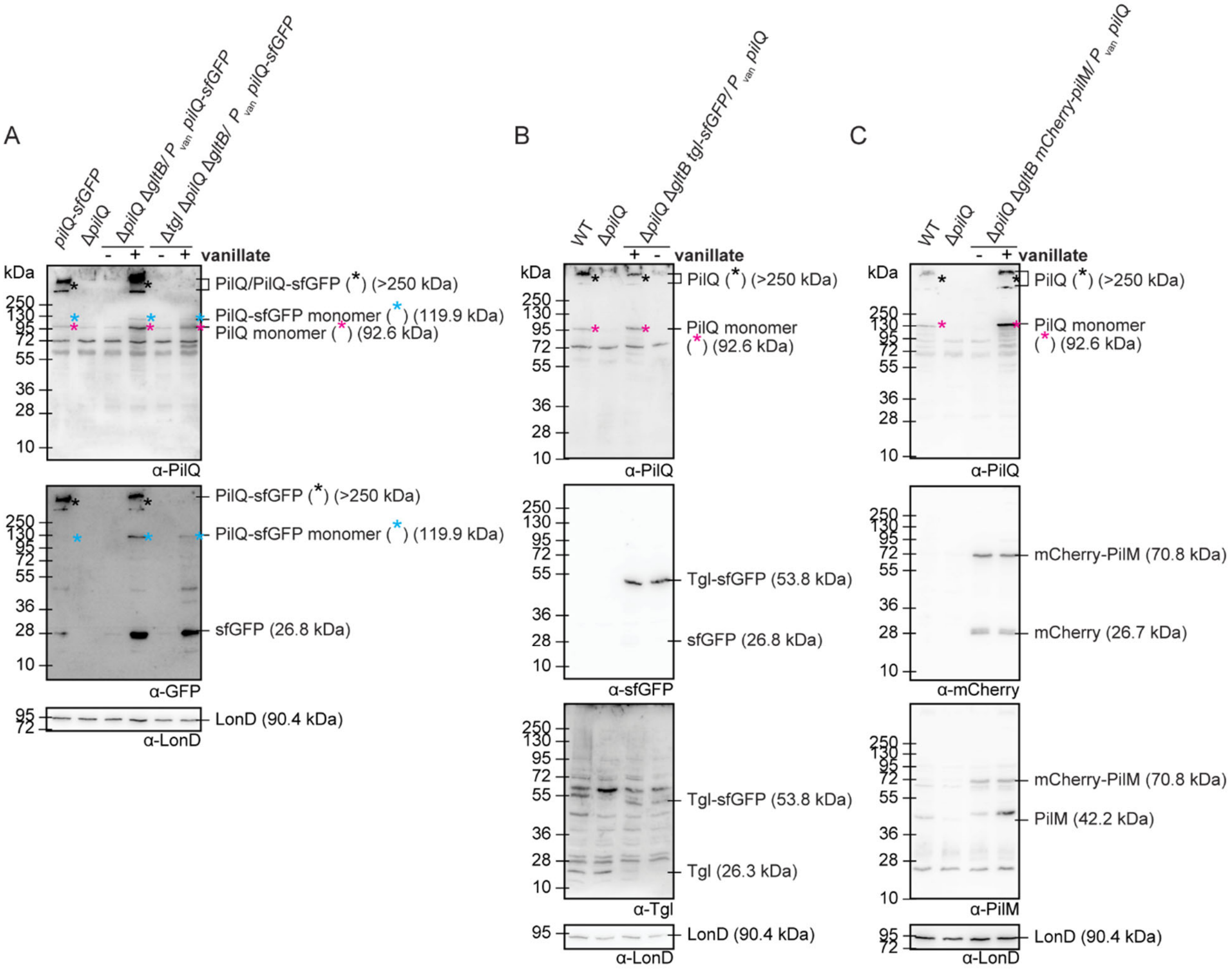


**Figure S2. Induction of PilQ synthesis**

(A-C) Immunoblot detection of (A) PilQ-sfGFP or (C, D) PilQ protein expression in the indicated strains. Protein from the same number of cells from exponentially growing suspension cultures was loaded per lane. To induce protein expression from $P_{van}$, cells were treated with vanillate for 24 h. In A, 10 µM vanillate was used for inducing PilQ-sfGFP accumulation at WT levels in the Δ*pilQ* background, and 500 µM vanillate was used to highly induce PilQ-sfGFP accumulation in the Δ*tgl* Δ*pilQ* background. In B, 20 µM vanillate was used for inducing PilQ accumulation at WT levels in Δ*pilQ* cells expressing *tgl-sfGFP*. In C, 1mM vanillate was used to rapidly induce *pilQ* expression in cells co-expressing *mCherry-pilM* (Fig. S4C). The blot was stripped before applying a new antibody. LonD served as a loading control. Monomeric or oligomeric forms of PilQ/PilQ-sfGFP are marked with an asterisk. Calculated molecular weights of proteins without signal peptide (if relevant) are indicated.

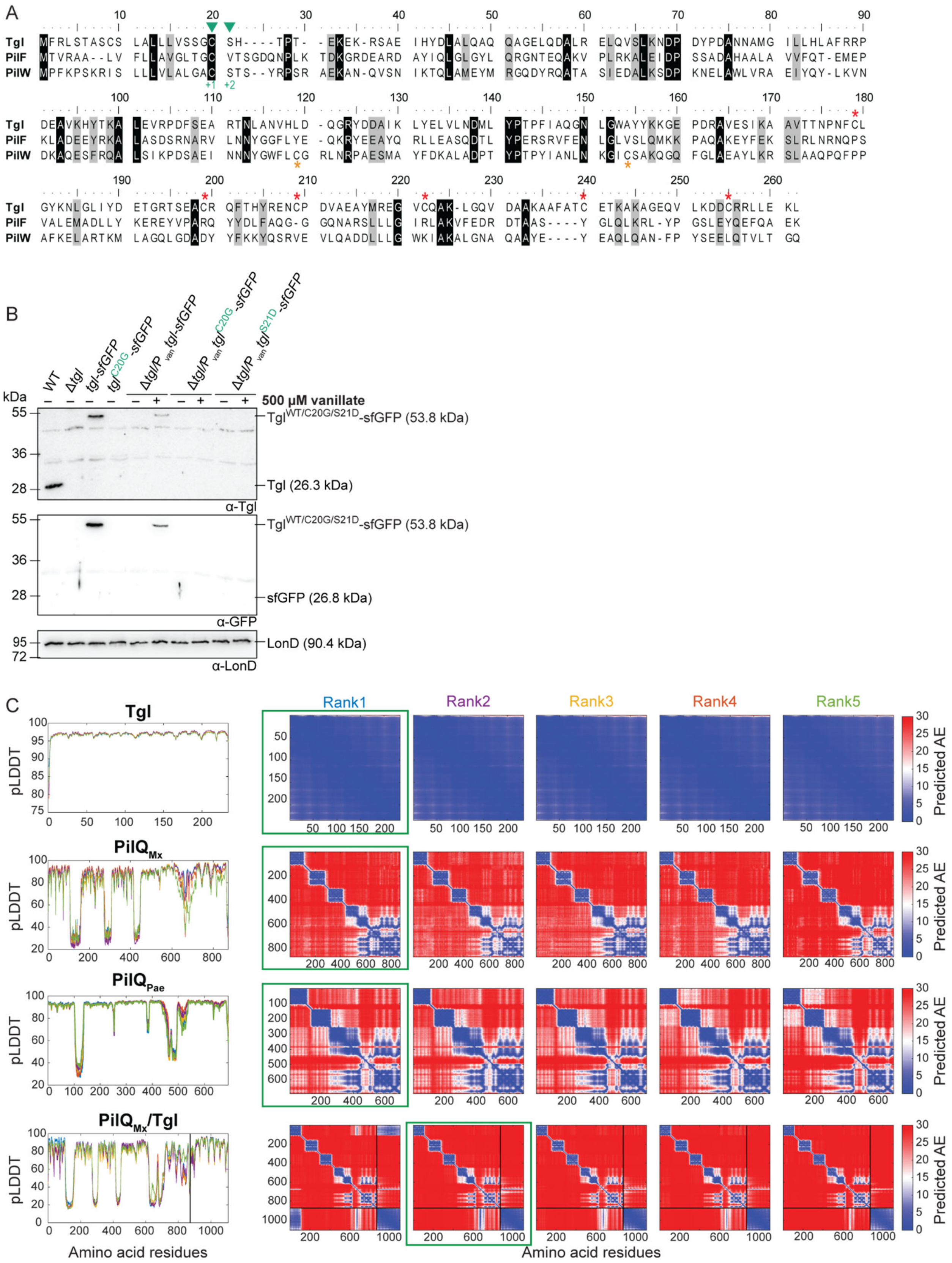

A
Tgl
PilF
PilW
+1 +2
B
WT
Δtgl
tgl-sfGFP
tgl$^{C20G}$-sfGFP
Δtgl/P$_{van}$ tgl-sfGFP
Δtgl/P$_{van}$ tgl$^{C20G}$-sfGFP
Δtgl/P$_{van}$ tgl$^{S21D}$-sfGFP
kDa
500 µM vanillate
Tgl$^{WT/C20G/S21D}$-sfGFP (53.8 kDa)
Tgl (26.3 kDa)
α-Tgl
sfGFP (26.8 kDa)
α-GFP
LonD (90.4 kDa)
α-LonD
C
Tgl
PilQ$_{Mx}$
PilQ$_{Pae}$
PilQ$_{Mx}$/Tgl
pLDDT
Rank1
Rank2
Rank3
Rank4
Rank5
Predicted AE
Amino acid residues

**Figure S3. Amino acid and structural characterization of TgI alone and in complex with its PilQ secretin partner, and accumulation of TgI-sfGFP variants**

(A) Alignment of full-length amino acid sequences of TgI, PilF from *P. aeruginosa,* and PilW from *N. meningitides*, where the acylated N-terminal Cys residue +1 of the mature lipoprotein and residue +2, important for retention of lipoproteins in the IM in *Escherichia coli*, are marked with a green arrow. The Cys residues within TgI predicted by AlphaFold to form disulfide bridges are marked with a red asterisk (Fig. 7A). The Cys residues within PilW that form a disulfide bridge are marked with an orange asterisk.

(B) Immunoblot detection of TgI variants. Protein from the same number of cells from exponentially growing suspension cultures was loaded per lane. To induce gene expression from $P_{van}$, cells were incubated with 500 μM vanillate for 24 h when indicated. Blot was probed with the indicated antibodies. The blot was stripped before applying a new antibody. LonD served as a loading control. Calculated molecular weights of proteins without signal peptide (if relevant) are indicated.

(C) pLDDT and pAE plots for five models of the indicated proteins or protein complexes in Fig. 7A, F and Fig. S5A, predicted by AlphaFold. The model marked by a green box was used for further analysis. Signal peptides were removed before generating a model.

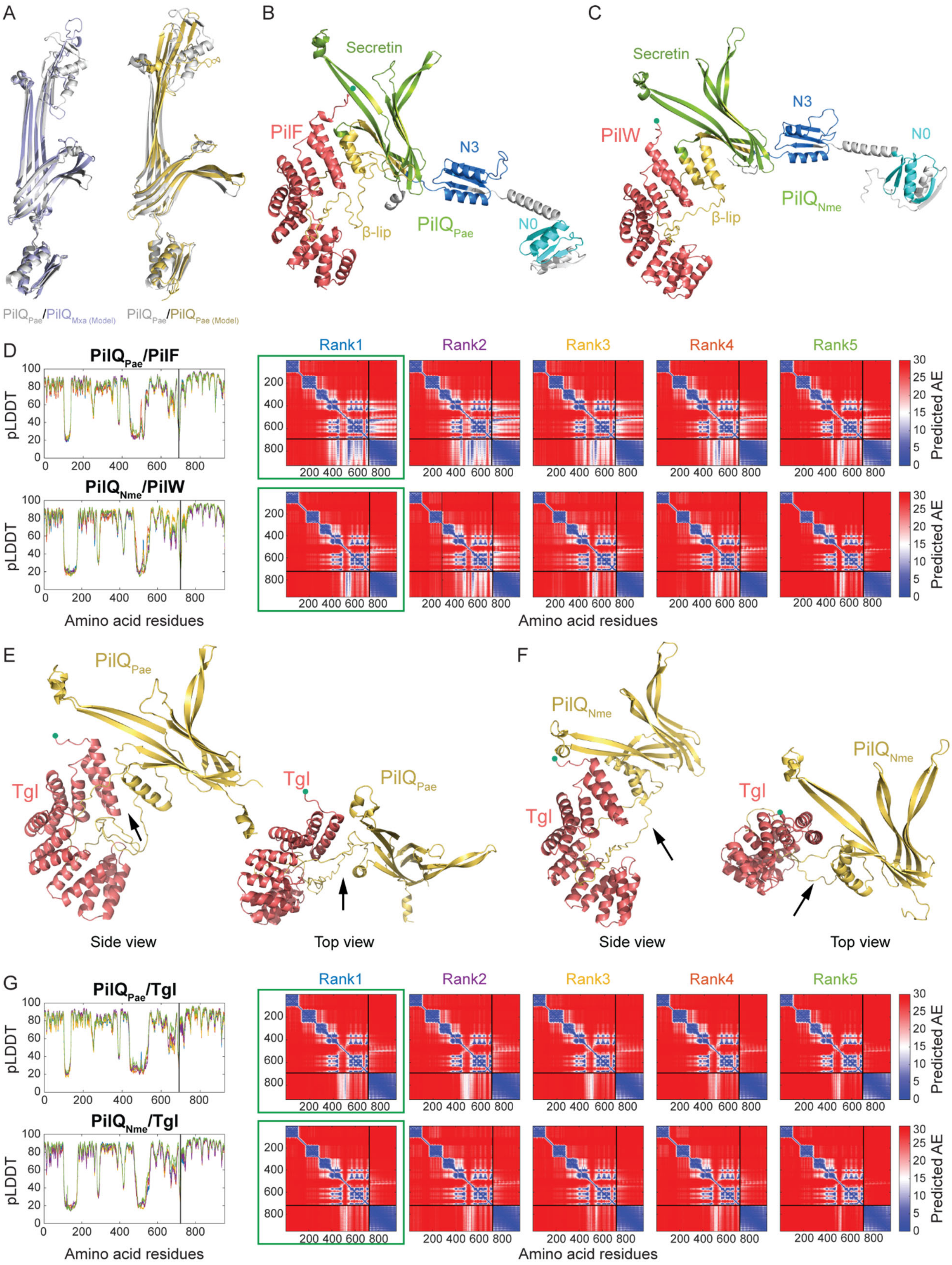
A
B
C
Secretin
PilF
N3
N0
β-lip
PilQPae
Secretin
PilW
N3
N0
β-lip
PilQNme
PilQPae/PilQMxa (Model)
PilQPae/PilQPae (Model)
D
PilQPae/PilF
PilQNme/PilW
pLDDT
Amino acid residues
Rank1
Rank2
Rank3
Rank4
Rank5
Predicted AE
Amino acid residues
E
F
PilQPae
Tgl
Tgl
PilQPae
PilQNme
Tgl
Tgl
Tgl
PilQNme
Side view
Top view
Side view
Top view
G
PilQPae/Tgl
PilQNme/Tgl
pLDDT
Amino acid residues
Rank1
Rank2
Rank3
Rank4
Rank5
Predicted AE
Amino acid residues

**Figure S4. Structural characterization of pilotin/secretin complexes**

(A) Superposition of the AlphaFold modeled PilQ structures of *M. xanthus* (lilac) or *P. aeruginosa* (yellow) with a PilQ protomer (gray) of the CryoEM structure of the secretin of *P. aeruginosa* in Fig. 7D.

(B, C, E, F) Modeled AlphaFold heterodimer of PilF/$PilQ_{Pae}$ (B), PilW/$PilQ_{Nme}$ (C), Tgl/$PilQ_{Pae}$ (E) and Tgl/$PilQ_{Nme}$ (F). The the acylated N-terminal Cys residue of mature Tgl (residue Cys20 in the unprocessed protein) that places the protein at the inner leaflet of the OM is indicated by a green circle. The domains of PilQ are colored as described in Fig. 7F.

(D, G) pLDDT and pAE plots for five models of the indicated proteins or protein complexes in panels B-C, E-F predicted by AlphaFold. The model marked by a green box was used for further analysis. Signal peptides were removed before generating a model.

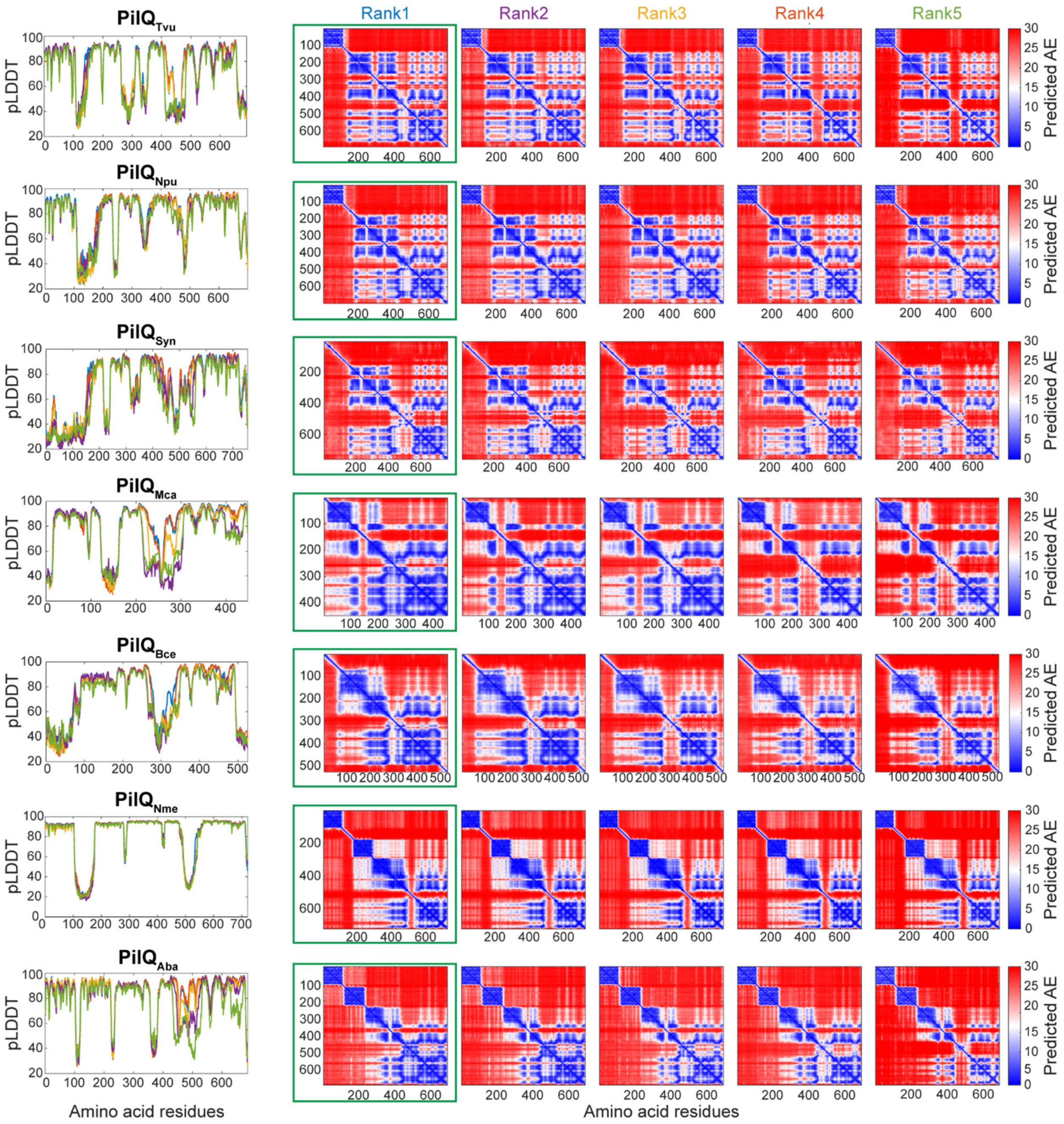


**Figure S5. pLDDT and pAE plots of the Alphafold models of PilQ secretins in other bacteria**

pLDDT and pAE plots for five models of the indicated proteins in Fig. 8 C-I predicted by AlphaFold. The model marked by a green box was used for further analysis. Signal peptides were removed before generating a model.

**Table S1.** Oligonucleotides used in this work[1]

| Primer name | Sequence 5'-3' | Brief description |
|---|---|---|
| tgl_fw_hindiii | ATCAAGCTTATGTTCCGCCTTTCCACCGCGTCC | For replacement of *tgl* with *tgl*-sfGFP |
| sfgfp_rv_xbaI | ATCTCTAGATTATTTGTAGAGCTCATCCATGCCAT | |
| tgl_ds_fw_xbaI | ATCTCTAGACGCGTTGGGTATCTGGGGACCCGT | |
| tgl_ds_rv | ATCGAATTCGGAGCGCCGCCACACAGCGAGCAC | |
| tgl_CtoG_A_HindIII | GCGCAAGCTTGTGATGATCGACAATCCCAT | For replacement of *tgl* with $tgl^{C20G}$ |
| tgl_CtoG_Bov | GGGCGTGTGGGAGCCACCGGAGGACACCAGCAGCA | |
| tgl_CtoG_Cov | GTGTCCTCCGGTGGCTCCCACACGCCCACGGAGAA | |
| tgl_CtoG_D_BamHI | GCGCGGATCCCCAGCTTTCGCCTTGGTCTCA | |
| Pvan_PilQ_fwd_NdeI | AGTCCATATGCTCGAGGAGAGCGCTGT | For expression of *pilQ or pilQ-sfGFP* from the *18-19* site under control of $P_{van}$ |
| sfGFP_rev_pilQ_EcoRI | TCAGGAATTCTTAGGATCCTTTGTAGAGC | |
| PilQ_rev_EcoRI | GCGTGAATTCTTACAGAGTCTGCGCAATGG | |
| Pvan_tgl_fw_NdeI | GATCCATATGTTCCGCCTTTCCACCGCG | For expression of tgl, *tgl-sfGFP*, $tgl^{C20G}$*-sfGFP or* $tgl^{S21D}$*-sfGFP* from the *18-19* site under control of $P_{van}$ |
| sfGFP_rv_tgl_EcoRI | CACTGAATTCTTATTTGTAGAGCTCATCCATGCCAT | |
| Pvan forw | TGGACTCTAGCCGACCGACTGAGACGC | |
| Tgl_S21G_overlay_rev | TGGGCGTGTGGTCGCAACCGGAGGACACCA | |
| Tgl_S21G_overlay_fwd | CTCCGGTTGCGACCACACGCCCACGGAGAA | |
| PilQ_dAMIN_A_XbaI | ATATTCTAGACGCTGCGTCGACCGCGGGCA | For deletion of the AMIN domains of PilQ |
| PilQ_dAMIN_B | TCCCACGGTAGCGGGCACCATGCACCCTGGCGCCC | |
| PilQ_dAMIN_C | GGTGCATGGTGCCCGCTACCGTGGGAAGCGCGTAT | |
| pilQ_dAMIN_D_HindIII | GAATAAGCTTCGCAGGTTGAGCTGAAGCGCCC | |
| tgl-A_XbaI | ATATTCTAGATACCGCGGGCTGCCCGCC | For Δ*tgl* |
| tgl-D_EcoRI | TGACGAATTCACGAGCCGGTCGGACTCG | |

| | | |
|---|---|---|
| PilQAMIN_A_KpnI | TATAGGTACCATGCAGACAAGGTTCGCCT | For generation of *pilQ*$^{\text{AMINs (1-475)}}$-*sfGFP* using the *pilQ::pilQ-sfGFP* strain |
| PilQAMIN_sfGFP_overlay_rev | ggagccgccgccgccGGCCTGCTGGGGCGCGCCCT | |
| PilQamin_sfGFP_overlay_fwd | GCGCCCCAGCAGGCCggcggcggcggctccatgag | |
| sfGFP_rev_pilQ_EcoRI | TCAGGAATTCTTAGGATCCTTTGTAGAGC | |

[1] Underlined sequences indicate restriction sites.
